# Obesity-induced astrocyte dysfunction impairs heterosynaptic plasticity in the orbitofrontal cortex

**DOI:** 10.1101/2020.05.01.073205

**Authors:** Benjamin K. Lau, Ciaran Murphy-Royal, Manpreet Kaur, Min Qiao, Jaideep S. Bains, Grant R. Gordon, Stephanie L. Borgland

## Abstract

Overconsumption of highly palatable, energy dense food is considered a key driver of the obesity pandemic. The orbitofrontal cortex (OFC) is critical for reward valuation of gustatory signals, yet how the OFC adapts to obesogenic diets is poorly understood. Here we show that extended access to a cafeteria diet impairs astrocyte glutamate clearance, which leads to a heterosynaptic depression of GABA transmission onto pyramidal neurons of the OFC. This decrease in GABA tone is due to an increase in extrasynaptic glutamate, which acts via metabotropic glutamate receptors to liberate endocannabinoids. This impaired the induction of endocannabinoid-mediated long-term plasticity. In obese rats, this cascade of synaptic impairments was rescued by restoring astrocyte glutamate transport with the nutritional supplement, N-acetylcysteine. Together, our findings indicate that obesity targets astrocytes to disrupt the delicate balance between excitatory and inhibitory transmission in the OFC.

**Highlights:**

- Diet-induced obesity induces hypertrophy of astrocytes and impairs their ability to transport glutamate.
- Failure of astrocytes to clear extrasynaptic glutamate drives endocannabinoid-mediated inhibitory long-term depression of principal output neurons in the OFC.
- Astrocytic glutamate transporter function is restored with NAC, which rescues the synaptic deficits.

## Introduction

Obesity is characterized by a disruption in energy balance, in which intake of energy far exceeds its output (Spiegelman and Flier, 2001). A key contributor to excess energy intake is overeating, which is especially pertinent in modern times given the abundance of palatable, hypercaloric food. Furthermore, once obesity is established it is highly persistent, such that weight loss is rarely maintained (Hall and Guo, 2017). A hypothesis underlying the persisting effects of obesity is that adaptive plasticity occurs in neural circuits, leading animals to defend elevated body weight (Matikainen-Ankney and Kravitz, 2018). Neural plasticity associated with feeding regulation, energy homeostasis and obesity has been well characterized within hypothalamic and mesolimbic systems (Matikainen-Ankney and Kravitz, 2018; Thoeni et al., 2020). However, there is growing evidence implicating cortical systems involved in decision-making and response inhibition (Lowe et al., 2019; Seabrook and Borgland, 2020). The orbitofrontal cortex (OFC) plays a key role in processing food-related signals (Jennings et al., 2019; Seabrook and Borgland, 2020). By integrating afferent and efferent projections from sensory, limbic and prelimbic regions (Ongür and Price, 2000), the OFC guides decision-making associated with food intake, such as processing value and updating actions based on current energy status (Seabrook and Borgland, 2020). However, less is known about how obesogenic diets influence plasticity in cortical regions.

One of the hallmarks of obesity is inflammation and astrocyte reactivity in the brain. Following short and long-term exposure to an obesogenic diet, astrocytes undergo morphological and functional changes leading to a hypertrophic and/or reactive phenotype associated with an inflammatory response (Douglass et al., 2017; García-Cáceres et al., 2013). However, the underlying mechanisms linking obesity-induced changes in astrocytes to neuronal plasticity are not well understood. Under physiological conditions, astrocytes provide metabolic support for neuronal synaptic function. Forming part of a tripartite synapse, astrocytes can regulate synaptic transmission through the uptake of neurotransmitters in the synaptic cleft (Chen et al., 2019; Hirase et al., 2014). In particular, astrocytes specifically express glutamate transporter 1 (GLT-1), responsible for the predominant clearance of the excitatory neurotransmitter, glutamate (Tanaka et al., 1997). While there is some evidence in other regions indicating obesity-induced changes in GLT-1 levels (Linehan et al., 2018; Tsai et al., 2018), the functional consequences of altered astrocytic glutamate transport on downstream neurotransmitter systems is lacking.

Using a diet known to produce reward dysfunction and overeating (Johnson and Kenny, 2010), we previously demonstrated that an obesogenic influences spine density and suppresses GABAergic synaptic transmission in the lateral OFC (lOFC) (Thompson et al., 2017). However, the mechanisms behind these synaptic changes and the impact on synaptic plasticity is not known. Here, we hypothesize that overnutrition with palatable food readily available for human consumption alters heterosynaptic plasticity in the OFC via a neuron-astrocyte interaction.

## Results

### An obesogenic diet alters inhibitory GABAergic synaptic transmission via recruitment of endocannabinoids

To determine whether diet-induced obesity alters synaptic function in the lOFC, we gave 24h (extended), 1h (restricted), or no access (chow) to a high fat, energy-dense cafeteria diet for 40 consecutive days, in addition to *ad libitum* access to chow (Figure 1A). Consistent with previous reports (Johnson and Kenny, 2010; Rolls et al., 1980; Thompson et al., 2017), rats given extended access to a cafeteria diet gained significantly more weight than those with restricted or chow access (Figure 1B). We have previously shown that this weight gain is associated with a higher energy intake, predominantly due to consumption of cafeteria diet rather than chow, and is associated with elevated plasma leptin and insulin levels consistent with obesity (Thompson et al., 2017). While restricted access rats exhibit similar preference and macronutrient intake of cafeteria diet, their weight gain was similar to chow rats as observed previously (Johnson and Kenny, 2010; Thompson et al., 2017). Thus, restricted access rats serve as a pair fed control whereby their ability to consume the diet in excess of that eaten by chow controls was prevented by the limited duration of exposure (Ellacott et al., 2010).

**Figure 1.**
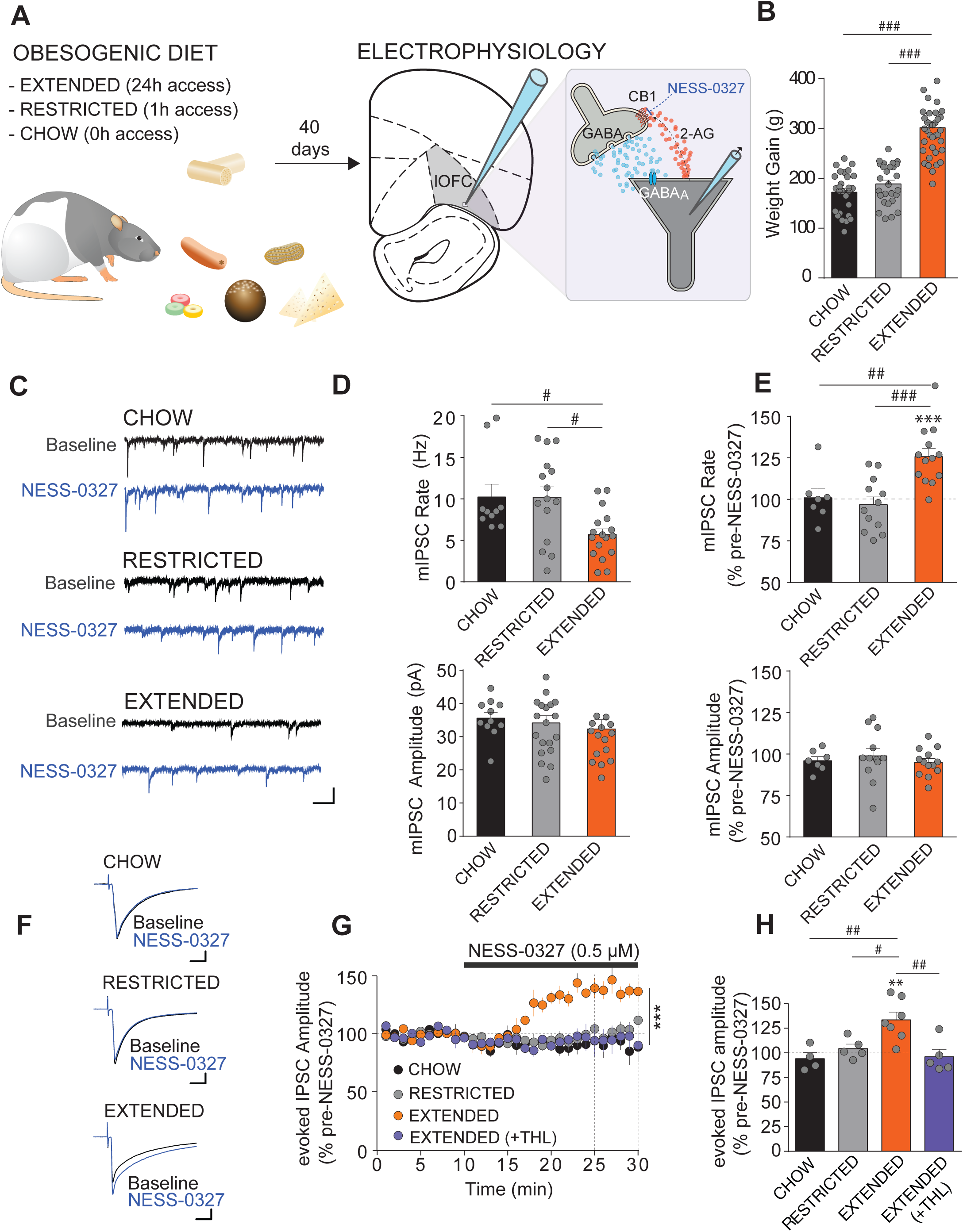
Extended access to a cafeteria diet leads to an endocannabinoid-mediated reduction in inhibitory GABAergic synaptic transmission onto lOFC pyramidal neurons. **(A)** Schematic of experimental paradigm: Rats were given 24 h (extended), 1 h (restricted) or 0 h (chow) access to a cafeteria diet for 40 days. In-vitro patch-clamp electrophysiology was then conducted in layer II/III pyramidal neurons from brain slices containing the lOFC. **(B)** Change in body weight following 40 days access to either chow (N = 27), restricted (1 h/day; N = 27), or extended (24 h/day, N = 38) access to a cafeteria diet. Rats with extended access to a cafeteria diet gain significantly more weight compared to those on chow (P < 0.001) or with restricted access (P < 0.001). One way ANOVA, F(2,89) = 68.49, P <0.001, Tukey’s post-hoc tests. **(C)** Example traces of mIPSCs before (Baseline) and during application of the neutral CB1 receptor antagonist, NESS-0327 (0.5 µM) in lOFC pyramidal neurons from chow, restricted or extended access rats. Scale bars: 50 pA, 100 ms. **(D)** Basal rate (upper) and amplitude (lower) of mIPSCs onto lOFC neurons from chow (n = 10/6), restricted (n = 14/7) or extended (n = 17/8) access rats. Rats with extended access to the cafeteria diet have decreased mIPSC rate compared to chow and restricted access rats (One-way ANOVA, F(2,38)=5.107, P = 0.011 with Tukey’s multiple comparison test). There was no significant difference in mIPSC amplitude between diet groups (F(2,38)=0.380, P = 0.686). **(E)** Percentage change in mIPSC rate (upper) and amplitude (lower) during application of NESS-0327 (0.5 µM) in lOFC pyramidal neurons from chow (n = 8/5), restricted (n = 12/5) or extended (n = 13/8) access rats. NESS-0327 unmasks a facilitation of mIPSC rate in extended access rats (P < 0.001^###^), which is significantly different from chow (P < 0.01^##^) and restricted (P < 0.001^###^) access rats (F(2, 30) = 11.37, P = 0.0002). NESS-0327 had no significant effect on amplitude (F(2,30) = 0.367, P= 0.696). mIPSC rate in lOFC neurons from extended access rats was significantly greater than baseline (t(12) = 5.326, p = 0.0002***), whereas mIPSC rate from chow (t(7) = 0.212, p = 0.838) and restricted access were not (t(11) = 0.682, p =0.509). **(F)** Example traces of evoked IPSC amplitude before (black) and during application of NESS-0327 (blue) in chow (top), restricted (middle), and extended (bottom) access rats. Scale bars: 100 pA, 10 ms. **(G)** Time course of evoked IPSC amplitude during application of NESS-0327 in lOFC pyramidal neurons from chow (n = 4/3), restricted (n = 5/4) and extended access (n = 7/4 without THL, n = 5/4 with THL) rats. Dotted vertical lines indicate the time period analyzed in (H). There was a significant interaction of diet group x timecourse (F(87,527) = 4.960, P <0.0001) and a main effect of diet (F(3,19) = 10.89, P=0.002 (Mixed-effect model 2-way RM ANOVA). **(H)** Bar graph quantifying the percentage change in evoked IPSC amplitude during application of NESS-0327 in lOFC pyramidal neurons from chow (n = 4/3), restricted (n = 5/4) or extended (n = 7/4 without THL (Diacylglycerol lipase inhibitor, tetrahydrolipstatin) n = 5/4 with THL) access rats. NESS-0327 unmasked a facilitation of evoked IPSC amplitude in lOFC pyramidal neurons from extended access rats (P < 0.01), which was blocked in the presence of an inhibitor of 2-AG synthesis, THL (P>0.05) (one way ANOVA: F(3, 17) = 7.445, P = 0.0021). Evoked IPSC amplitude in lOFC neurons from extended access rats was significantly greater than pre-NESS-0327 baseline (t(6) = 4.246, p = 0.0054**, whereas evoked IPSCs from lOFC neurons of chow (t(3) = 0.9659, P =0.4054), restricted access (t(4) = 0.9428, P = 0.3991) or extended access + THL (t(4) = 0.6231, P = 0.6231) were not. Bars represent mean ± s.e.m with individual values overlaid.

To compare basal GABAergic input to lOFC neurons between the dietary conditions, miniature inhibitory postsynaptic currents (mIPSCs) were recorded in layer II/III pyramidal neurons from extended, restricted and chow access rats (Figure 1A). We found that the rate, but not amplitude of mIPSCs onto lOFC pyramidal neurons was significantly decreased in extended access rats compared to chow or restricted access rats (Figure 1C,D), consistent with a presynaptic mechanism of action reported previously (Thompson et al., 2017). To test whether these changes in GABA transmission were specific to the lOFC, we additionally examined mIPSCs in adjacent cortical regions, including the prelimbic cortex and primary motor cortex (M1), located dorsomedial and dorsal of the lOFC, respectively (Figure S1A). In contrast to the lOFC, we found that the rate and amplitude of mIPSCs were not significantly different between chow and extended access rats in both the prelimbic and motor cortex (Figure S1B-D).

Obesity is associated with elevated levels of the major endocannabinoids, 2-arachidonoylglycerol (2-AG) and anandamide (AEA) in both the brain and periphery (Gatta-Cherifi and Cota, 2016). This is associated with impairments in endocannabinoid signalling and plasticity (Cristino et al., 2013; Linehan et al., 2018; Massa et al., 2010). We hypothesized that elevated levels of endocannabinoids may mediate the suppression of inhibition observed in the lOFC of extended access rats. If endocannabinoids are tonically present under basal conditions to suppress GABAergic transmission, then blockade of CB1 receptors should unmask a facilitation of inhibitory transmission. To test this idea, we examined the effect of the neutral CB1 receptor antagonist, NESS-0327 (0.5 μM) on GABA release events. We chose a neutral CB1 receptor antagonist as this would preclude effects due to constitutive receptor activity, unlike the prototypical CB1 receptor inverse agonist, AM251. Indeed, NESS-0327 (0.5 μM) facilitated the rate of mIPSCs in lOFC pyramidal neurons from extended access rats (126 ± 5%), but did not change that of restricted (96 ± 5%), or chow (101 ± 6%) access rats (Figure 1E). NESS-0327 had no effect on the amplitude of mIPSCs in lOFC neurons from chow (96 ± 2%), restricted (99 ± 5%) or extended (95 ± 2%) access rats (Figure 1E). Together, this indicates the presence of a basal endocannabinoid tone suppressing GABA release onto lOFC neurons from extended, but not chow or restricted access rats. To confirm that this effect can also occur with action potential-driven activity, we repeated the experiment with electrically-evoked IPSCs. Similarly, NESS-0327 (0.5 µM) application unmasked a facilitation of evoked IPSCs in lOFC neurons from extended access (133 ± 8%), but not chow (94 ± 6%) or restricted access rats (104 ± 5%, Figure 1F-H). The facilitation of evoked IPSC amplitude by NESS-0327 was abolished in slices pre-incubated with the diacyglycerol lipase inhibitor, tetrahydrolipstatin (THL, 10 μM, 96 ± 7%, Figure 1G-H), which prevents the biosynthesis of the endocannabinoid 2-AG. This indicates the presence of a 2-AG-mediated endocannabinoid tone suppressing evoked GABAergic transmission onto lOFC pyramidal neurons in obese rats. We additionally examined whether endocannabinoid tone modulates glutamatergic synapses. Application of the CB1 agonist, WIN55,212-2 (1 µM) in chow access rats produced inhibition of evoked excitatory postsynaptic currents (EPSCs) onto lOFC pyramidal neurons (77 ± 4%, Figure S2A-C), indicating that glutamatergic synapses are sensitive to endocannabinoids. However, application of NESS-0327 (0.5 µM) in extended access rats had no effect on evoked EPSCs (98 ± 6%, Figure S2D-F). Together, these data suggest that while both excitatory and inhibitory synapses onto lOFC pyramidal neurons are sensitive to endocannabinoids, only inhibitory synapses exhibit a functional endocannabinoid tone during the obese state.

### Endocannabinoid-mediated long-term depression is impaired in the lOFC of obese rats

We next investigated if the presence of an endocannabinoid tone in the lOFC of obese rats was associated with impairments in synaptic plasticity. We first tested if inhibitory synapses onto lOFC pyramidal neurons undergo long-term depression (LTD) in diet-naive rats (Figure S3A). We found that strong theta-burst stimulation (TBS), mimicking endogenous high-frequency gamma bursts at a lower-frequency theta oscillation (Buzsáki and Wang, 2012), induced a long-lasting suppression of evoked IPSC amplitude onto lOFC pyramidal neurons (Figure S3B-D). This was associated with paired-pulse facilitation (Figure S3E), suggesting a presynaptic mechanism of action. This presynaptic, TBS-induced LTD was consistent with endocannabinoid-mediated LTD reported in other brain regions (Chevaleyre and Castillo, 2004; Jiang et al., 2010; Zhao et al., 2015). In particular, previous work in the visual (Jiang et al., 2010) and somatosensory (Zhao et al., 2015) cortex has demonstrated that TBS-LTD specifically requires endocannabinoids. Therefore, we tested if the CB1 receptor antagonist, AM251 could block TBS-induced LTD in lOFC neurons. TBS in the presence of AM251 (3 µM) blocked LTD and induced a small facilitation of evoked IPSC amplitude (Figure S3B-D). Given that many forms of endocannabinoid-mediated LTD depend on activation of extrasynaptic group 1 metabotropic glutamate receptors (mGluR1/5), we further examined TBS-LTD in the presence of the mGluR5 antagonist, MTEP. MTEP also abolished TBS-LTD of evoked IPSC amplitude of lOFC neurons (Figure S3B-D), indicating that this form of LTD requires activation of mGluR5 in addition to CB1 receptors. Confirming this, AM251 or MTEP inhibited the paired-pulse facilitation induced by TBS (Figure S3E). Together, TBS can induce LTD of lOFC GABA synapses that is dependent on activation of mGluR5 and endocannabinoids.

Having established that inhibitory synapses in the lOFC can undergo endocannabinoid-mediated plasticity, we next examined if TBS-LTD was altered by diet-induced obesity. Following TBS, a long-term suppression of evoked IPSC amplitude was observed in lOFC pyramidal neurons of chow and restricted access rats (69 ± 9% and 67 ± 5% of baseline, respectively, Figure 2A-B), and this was associated with an increase in paired-pulse ratio in chow access rats (Figure 2C). By contrast, TBS had no effect on evoked IPSCs in lOFC pyramidal neurons of extended access rats (98 ± 4% of baseline, Figure 2A-C). Taken together, endocannabinoid-mediated LTD is impaired in the lOFC of obese rats.

**Figure 2.**
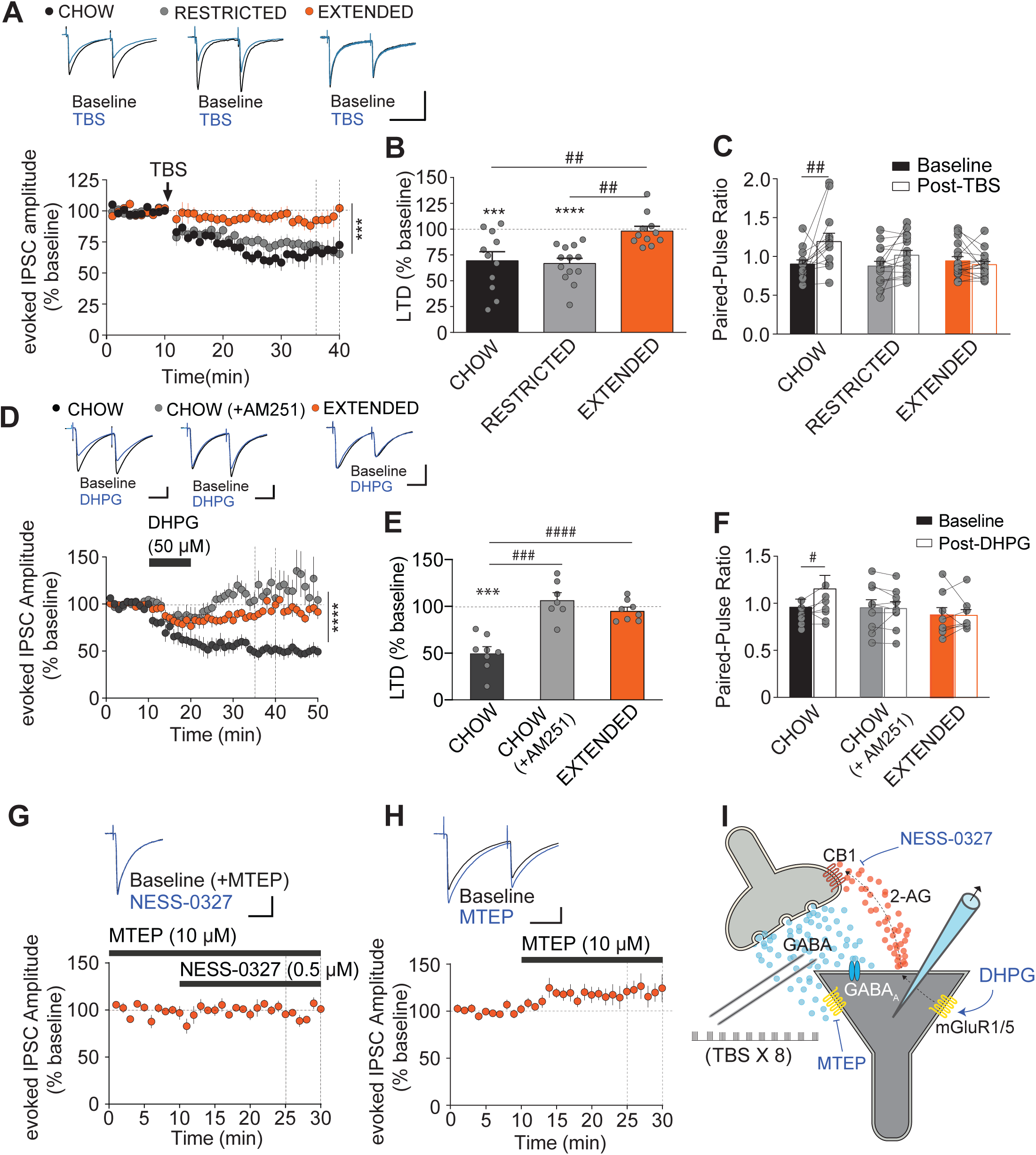
Endocannabinoid-mediated inhibitory LTD of lOFC pyramidal neurons is impaired with extended, but not chow or restricted access to a cafeteria diet. **(A)** Time plot of evoked IPSC amplitude before and after theta-burst stimulation (TBS) in lOFC neurons from chow (n = 11/9), restricted (n = 13/9) or extended (n = 11/7) access rats. Dotted vertical lines indicate the 25-30 min time period following TBS analyzed in B. There was a significant interaction of diet group x timecourse (F(116,2053) = 2.819, P <0.0001) and a main effect of diet (F(2,44) = 11.00, P=0.0001*** (Mixed-effect model 2-way RM ANOVA). Inset: Example traces of evoked IPSC amplitude before and after TBS in lOFC pyramidal neurons from chow, restricted or extended access rats. Scale bars: 200 pA, 20 ms. **(B)** Bar graph quantifying the percentage change in evoked IPSC amplitude 25-30 min following TBS in lOFC neurons from chow, restricted or extended access rats. TBS-LTD is inducible in lOFC neurons of chow (P < 0.01^##^) and restricted (P < 0.01^##^) access rats, but is impaired in extended access rats (F(2,32)=7.511, P = 0.002). Evoked IPSCs after TBS is significantly decreased from baseline in lOFC neurons from chow (n = 11/9, t(10) = 3.495, P=0.0058***) and restricted access rats (n = 13/9, t(12) = 6.637, P<0.0001****), but not extended (n = 11/7, t(10) = 0.3991, P=0.6982) access rats. **(C)** Bar graph of the paired-pulse ratio of evoked IPSC amplitude before (Baseline) and after TBS (Post-TBS) in lOFC neurons from chow, restricted and extended access rats. A RM 2-way ANOVA indicates a significant diet x LTD interaction (F(2, 43) = 5.087, P = 0.010) with a main effect of TBS (F(1, 43) = 8.537, P = 0.0055). A Sidak’s multiple comparison test reveals a significant effect of TBS on paired-pulse facilitation in lOFC neurons from chow (n = 11/9, P < 0.01), but not restricted (n = 13/9) or extended access rats (n = 11/7). Bars represent mean ± s.e.m with individual values overlaid. **(D)** Time plot of evoked IPSC amplitude in lOFC neurons before, during and after the mGluR1/5 agonist, DHPG (50 µM) application in chow, restricted and extended access rats. DHPG induces an LTD in chow (n = 10/5) access rats, which is blocked in slices pre-treated with the CB1 receptor antagonist, AM251 (3 µM, n = 15/5). In contrast, DHPG does not induce LTD in extended (n = 12/6) access rats. Dotted vertical lines indicate the 15-20 min time period following DHPG application analyzed in E. A Mixed-effect model 2-way RM ANOVA indicates a significant diet group x LTD interaction (F(78,1140) = 4.684, P <0.0001) with a main effect of diet (F(2,34) = 12.85, P<0.0001) and a main effect of LTD (F(5.110,149.4) = 11.59, P <0.0001). Inset: Example traces of evoked IPSC amplitude before and after application of DHPG (50 µM) in slices from chow, chow + AM251 (3 µM) treated and extended access rats. Scale bars: 250 pA, 25 ms. **(E)** Bar graph quantifying the percentage inhibition of evoked IPSC amplitude by DHPG (15-20 min post-application) in lOFC pyramidal neurons from chow (n = 8/5), chow + AM251 (3 µM, n = 7/5), or extended access rats (n = 8/6). DHPG produces a long-term suppression of evoked IPSC amplitude, which is blocked in the presence of AM251 and absent in extended access rats (F(2,20) = 23.70, P < 0.0001, Tukey’s multiple comparison test: chow vs. chow + AM251, P<0.001^###^, chow vs. extended, P<0.0001^###^). Evoked IPSC amplitude after DHPG is significantly decreased from baseline in lOFC neurons from chow (n = 8/5, t(7) = 7.37, P=0.0002***), but not in the presence of AM251 (n = 7/5, t(6) = 0.923, P = 0.392) nor in extended access rats (n = 8/6, t(7) = 1.18, P=0.276). Cells were excluded if the recording did not last until the analysis period. **(F)** Bar graph of paired-pulse ratio of evoked IPSCs before and 15-20 min after application of DHPG in lOFC pyramidal neurons from chow (n = 9/5), chow + AM251 (n =9/5) and extended access rats (n = 8/6). A 2-way RM ANOVA: Diet x drug interaction: (F(2,23) = 2.690, P = 0.089). A Sidak’s multiple comparisons test indicates that DHPG elicits paired-pulse facilitation in lOFC neurons from chow (P<0.05^#^), but not in the presence of AM251, or in extended access rats. **(G)** The facilitation of evoked IPSC amplitude by the neutral CB1 receptor antagonist, NESS-0327 (0.5 µM) is blocked by the mGluR5 antagonist, MTEP in lOFC neurons from extended access rats (n = 7/5). Inset: Example traces of evoked IPSC amplitude before and during application of NESS-0327 from slices pre-incubated with MTEP in lOFC neurons from extended access rats. Scale bars: 250 pA, 25 ms. **(H)** In extended access rats (n = 7/5), MTEP unmasks a facilitation of evoked IPSC amplitude, indicative of an mGluR5-mediated tone onto lOFC pyramidal neurons. Bars represent mean ± s.e.m with individual values overlaid. Inset: Example traces of evoked IPSC amplitude before and during application of MTEP. Scale bars: 250 pA, 25 ms. (**I**) Experimental schematic illustrating (i) endocannabinoid-mediated inhibitory LTD in OFC pyramidal neurons induced via TBS. (ii) endocannabinoid-mediated inhibitory LTD of OFC pyramidal neurons induced via Group 1 mGluR activation with DHPG (50 µM, 10 mins). (iii) unmasking of endocannabinoid tone with the CB1 receptor antagonist, NESS-0327 (0.5 µM). This endocannabinoid tone is prevented by pre-treatment with the mGluR5 antagonist, MTEP (10 µM) and mimicked by MTEP application alone.

Given the link between Group 1 mGluR and endocannabinoid signaling, we next investigated whether the obesity-induced endocannabinoid tone and impairment of TBS-LTD was mediated upstream by activation of mGluR1/5. Similar to TBS-LTD, the mGluR1/5 agonist, DHPG (50 μM) produced an LTD of evoked IPSC amplitude in lOFC neurons from chow access rats, which was blocked in the presence of AM251 (Figure 2D-E). Consistent with a presynaptic effect, DHPG produced a paired-pulse facilitation, which was blocked by AM251 (Figure 2F). By contrast, in rats with extended access to a cafeteria diet, DHPG (50 µM) had no effect on evoked IPSCs onto lOFC pyramidal neurons (Figure 2D-F). Together, these results indicate that similar to TBS, mGluR1/5 activation induces endocannabinoid-mediated LTD. This plasticity is also impaired in obese rats, suggesting a disruption in mGluR1/5-endocannabinoid signaling.

We next tested if the endocannabinoid tone previously revealed by a neutral CB1 receptor antagonist in obese rats could be blocked with an mGluR5 antagonist. In the presence of MTEP, NESS-0327 (0.5 μM) no longer produced a facilitation of evoked IPSC amplitude in lOFC neurons from extended access rats (Figure 2G), indicating the necessity of mGluR5 activation in mediating endocannabinoid tone. Furthermore, MTEP application alone mimicked the facilitation of evoked IPSCs observed with NESS-0327 (Figure 2H). Together, these results indicate that mGluR5 activation induces endocannabinoid tone, which may occlude the induction of endocannabinoid-mediated LTD in the lOFC of obese rats (Figure 2I).

### An obesogenic diet induces an extrasynaptic glutamate tone in the OFC

The obesity-induced alteration in Group 1 mGluR signalling may be due to a change in receptor sensitivity, number, or an occlusion by enhanced levels of synaptic glutamate spill over. To address this, we first examined basal glutamatergic synaptic input onto lOFC neurons from chow or extended access rats (Figure 3A). In the presence of picrotoxin (100 μM) and TTX (0.5 μM), miniature excitatory postsynaptic currents (mEPSC) were readily observed (Figure 3B). While we observed no change in mEPSC amplitude between diet groups (Figure 3B,D), there was a significant reduction in the rate of mEPSCs onto lOFC neurons from extended access rats compared to chow access rats (Figure 3B,C), indicative of a decrease in presynaptic glutamate release. We additionally gauged synaptic glutamate levels by examining the effect of the low-affinity AMPA receptor antagonist, γ-DGG (1 µM) on evoked AMPA EPSCs (Figure 3E). γ-DGG competes with ambient glutamate to bind synaptic AMPA receptors. Thus, the inhibition of evoked AMPA EPSC amplitude produced by γ-DGG will inversely correlate with synaptic levels of glutamate. Compared to lOFC neurons from chow access rats, γ-DGG produced significantly greater inhibition of evoked AMPA EPSC amplitude in lOFC neurons from extended access rats (Figure 3F,G,I), indicating decreased synaptic glutamate. To interrogate if this effect was due to an alteration in synaptic glutamate or a change in AMPA receptor number or function, we applied the high-affinity AMPA receptor antagonist, DNQX (1 µM), which fully displaces ambient glutamate from binding to synaptic AMPA receptors. There was no difference in DNQX-induced inhibition of evoked AMPA EPSCs in lOFC neurons from chow or extended access rats (Figure 3H,I), which together with the pattern of γ-DGG inhibition suggests a lack of change in AMPA receptor number/function. Contrary to our prediction that synaptic glutamate spillover leads to mGluR5 activation, the above results suggest there is decreased synaptic glutamate release onto lOFC neurons from obese rats.

**Figure 3:**
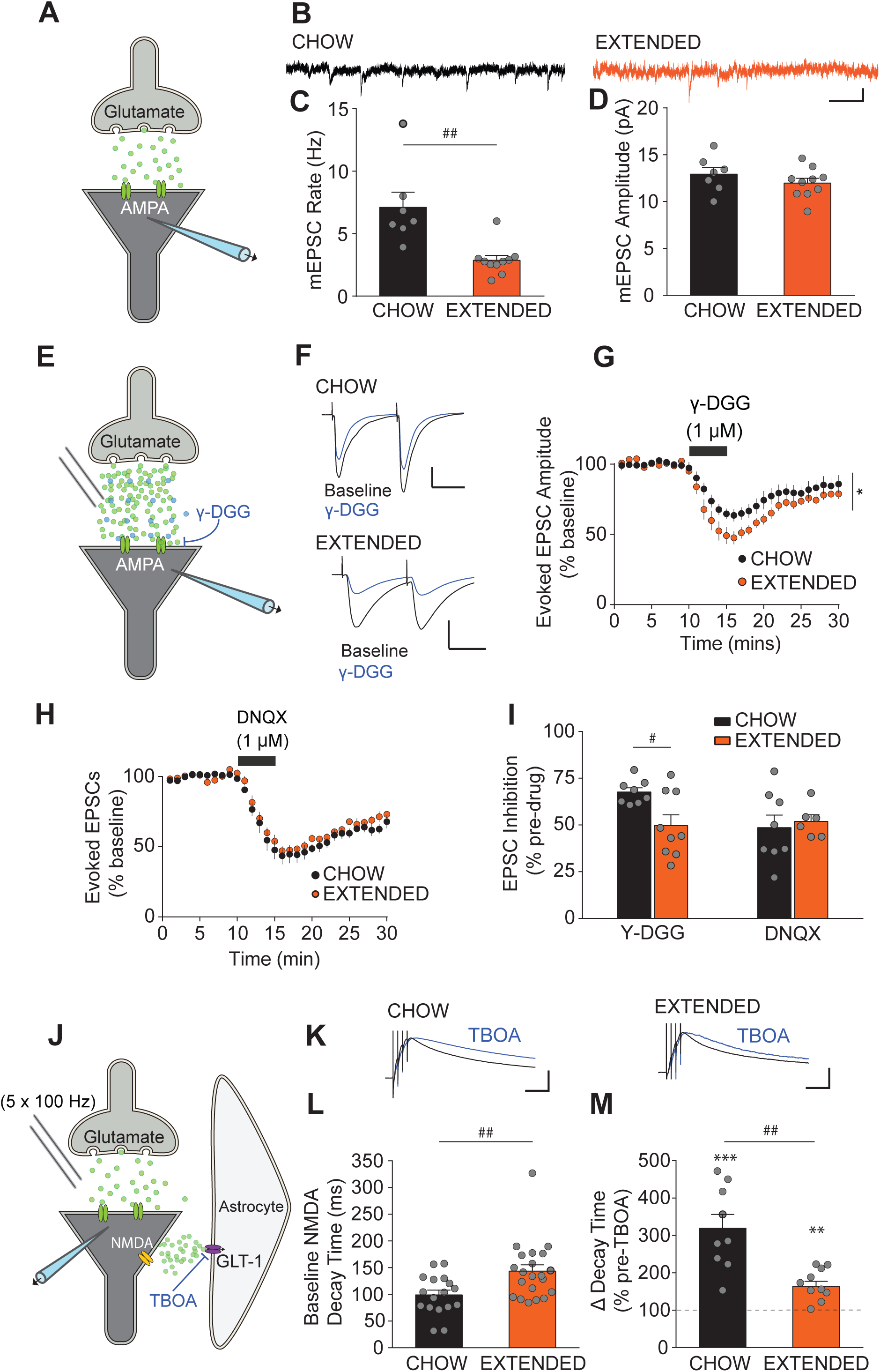
An obesogenic diet alters extrasynaptic and synaptic glutamate via impairment of astrocytic GLT-1. **(A)** Experimental schematic illustrating the recording of miniature excitatory postsynaptic currents (mEPSCs) in the presence of picrotoxin (100 μM) and tetrodotoxin (TTX, 0.5 μM). **(B)** Example traces of mEPSCs in lOFC pyramidal neurons from chow and extended access rats. Scale bars: 10pA, 0.1s. **(C)** Frequency of mEPSCs in lOFC pyramidal neurons from extended (n = 10/5) access rats is significantly reduced compared to that of chow (n = 7/4) access rats (t_(15)_=3.82, P = 0.0017^##^). Inset, **(D)** Amplitude of mEPSC is not different between lOFC pyramidal neurons of extended (n = 10/5) and chow (n = 7/4) access rats (t_(15)_=1.11, P = 0.285). **(E)** Experimental schematic illustrating the effect of the low affinity, competitive AMPA receptor antagonist, γ-DGG on evoked AMPA EPSCs. γ-DGG competes with glutamate to bind with AMPA receptors. Thus, the degree of AMPA EPSC inhibition produced by γ-DGG is inversely correlated with synaptic glutamate levels. **(F)** Example traces of evoked AMPA EPSCs before and during application of γ-DGG (1 μM) in chow (scale bar: 500 pA, 25 ms) and extended access rats (scale bar: 250 pA, 25 ms). **(G)** Time plot of evoked AMPA EPSC amplitude before and during γ-DGG (1 μM) application in lOFC pyramidal neurons from chow (n = 8/5) and extended (n/N = 9/7) access rats. A mixed-effects model (REML) reveals a significant diet x time interaction (F(29,422) = 3.135, P< 0.0001), main effect of diet (F(1,15) = 7.683, P = 0.0142*). **(H)** Time plot evoked of evoked AMPA EPSC amplitude before and during application of the high-affinity AMPA receptor antagonist, DNQX (10 μM) in lOFC neurons from chow (n = 8/5) or extended (n = 5/4) access rats. A mixed-effects model (REML) reveals a no diet x time interaction (F(29,290) = 0.531, P= 0.979), or main effect of diet (F(1,11) = 1.130, P = 0.310). **(I)** Bar graph quantifying the percentage inhibition of AMPA EPSC amplitude in response to γ-DGG and DNQX in lOFC pyramidal neurons from chow or extended access rats. A 2-way ANOVA indicates a significant interaction of diet x drug (F(1,26) = 4.45, P = 0.447). A Sidak’s multiple comparison test indicates that γ-DGG produces differential inhibition of evoked EPSC amplitude in lOFC neurons of chow compared to extended access rats (P < 0.05^#^). **(J)** Experimental schematic illustrating the measurement of extrasynaptic glutamate via extrasynaptically-located NMDA receptors. Evoked NMDA EPSCs are isolated by recording at +40 mV in the presence of DNQX (10 μM), picrotoxin (100 μM) and strychnine (1 μM). A high-frequency stimulation train (5 pulses @100 Hz) is used to elicit excess glutamate spillover into the extrasynaptic space, which is reflected as a prolongation in the decay time of evoked NMDA EPSCs. The effect of the glutamate transporter blocker, TBOA (30 μM) was then examined on this measure. **(K)** Example traces of evoked NMDA EPSCs in response to a 5 × 100 Hz stimulus normalized to peak amplitude before and during application of the glutamate transport blocker, TBOA in lOFC neurons from chow or extended access rats. Stimulus artifacts have been minimized for clarity. Scale bars: 0.5, 50 ms. **(L)** Half-decay time of evoked NMDA EPSCs during an initial baseline period (1 min) immediately following whole-cell access to a lOFC pyramidal neuron. Half-decay time is significantly longer in lOFC pyramidal neurons of extended (n/N = 21/5) access rats compared to chow (n/N = 17/6) rats (Unpaired t-test, t_(36)_ = 2.89, P = 0.0065^##^). **(M)** Change in decay time of evoked NMDA EPSCs in response to 10 min application of TBOA. Decay time is enhanced by TBOA in lOFC neurons of both chow (n = 9/4, t_(8)_=5.956, P = 0.0003***) and extended access rats (n = 9/6, t_(8)_=4.885, P = 0.0012**) compared to baseline. However, this enhancement is significantly impaired in lOFC pyramidal neurons from extended access rats (t_(16)_=3.97, P = 0.0011^##^). Bars represent mean ± s.e.m with individual values overlaid.

Given that obese rats have decreased synaptic glutamate release onto OFC pyramidal neurons, we explored an alternate hypothesis that Group 1 mGluRs are activated by enhanced levels of extrasynaptic glutamate. First, to indirectly interrogate levels of extrasynaptic glutamate, we induced spillover of glutamate onto extrasynaptic NMDA receptors under conditions of high activity (Figure 3J). Thus, NMDA receptors served as sensor for extrasynaptic glutamate. Pharmacologically isolated NMDA EPSCs were evoked at +40 mV in response to a high frequency train of electrical stimulation (5 x 100 Hz) (Figure 3K). To measure the duration of glutamate action at NMDA receptors, the half-decay time of evoked NMDA EPSCs was analyzed. We found that baseline decay time was significantly prolonged in lOFC neurons from extended access rats relative to chow access rats (Figure 3K,L), indicating elevated levels of glutamate in the extrasynaptic space. To indirectly assess changes in the efficacy of glutamate transport, we examined the action of the non-selective glutamate transport inhibitor, DL-TBOA (TBOA). In chow access rats, TBOA (30 μM) produced a 319 ± 37% increase in decay time, but this facilitation was significantly reduced to 164 ± 13% in lOFC neurons from extended access rats (Figure 3K,M). TBOA was less effective at preventing extrasynaptic glutamate reuptake in extended access rats, suggesting that there is a partial occlusion of NMDA receptors by a tonic presence of extrasynaptic glutamate. Thus, extrasynaptic glutamate tone was likely the result of reduced efficacy of non-synaptic glutamate transporters, rather than synaptic spillover of glutamate.

Given the observed increase in extrasynaptic glutamate but decrease in synaptic glutamate, we next tested if this compartmentalization of glutamate was mechanistically linked. One possibility for decreased synaptic glutamate release is through autoinhibition via presynaptic Group II mGluR (mGluR2/3) activation by extrasynaptic glutamate. To test this possibility, we examined the effect of the mGluR2/3 agonist, LY379268 on evoked AMPA EPSCs in lOFC neurons from chow or extended access rats (Figure 4A). LY379268 (100 nM) inhibited evoked EPSCs from chow access rats, which was reversed following addition of the mGluR2/3 antagonist, LY341495 (1 μM)(Figure 4B-D). Thus, glutamatergic synapses of OFC pyramidal neurons are sensitive to extrasynaptic inhibition via mGluR2/3 activation. Compared to chow access rats, the LY379268 agonist produced significantly less inhibition of EPSCs in lOFC neurons from extended access rats (Figure 4B-D), indicating possible occlusion of mGluR2/3 by an extrasynaptic glutamate tone. To test this, we examined the effect of the LY341495 antagonist alone on evoked AMPA EPSCs in lOFC neurons from chow or extended access rats (Figure 4E). If an extrasynaptic glutamate tone is suppressing excitatory transmission under basal conditions, an mGluR2/3 antagonist should unmask a facilitation of this response. LY341495 (1 μM) had no effect on evoked EPSCs in lOFC neurons from chow access rats, but significantly facilitated this response in extended access rats (Figure 4F-H). Together, these data suggest that in obese rats, there is a persistent suppression of synaptic glutamate release via tonic activation of presynaptic mGluR2/3 receptors by extrasynaptic glutamate.

**Figure 4:**
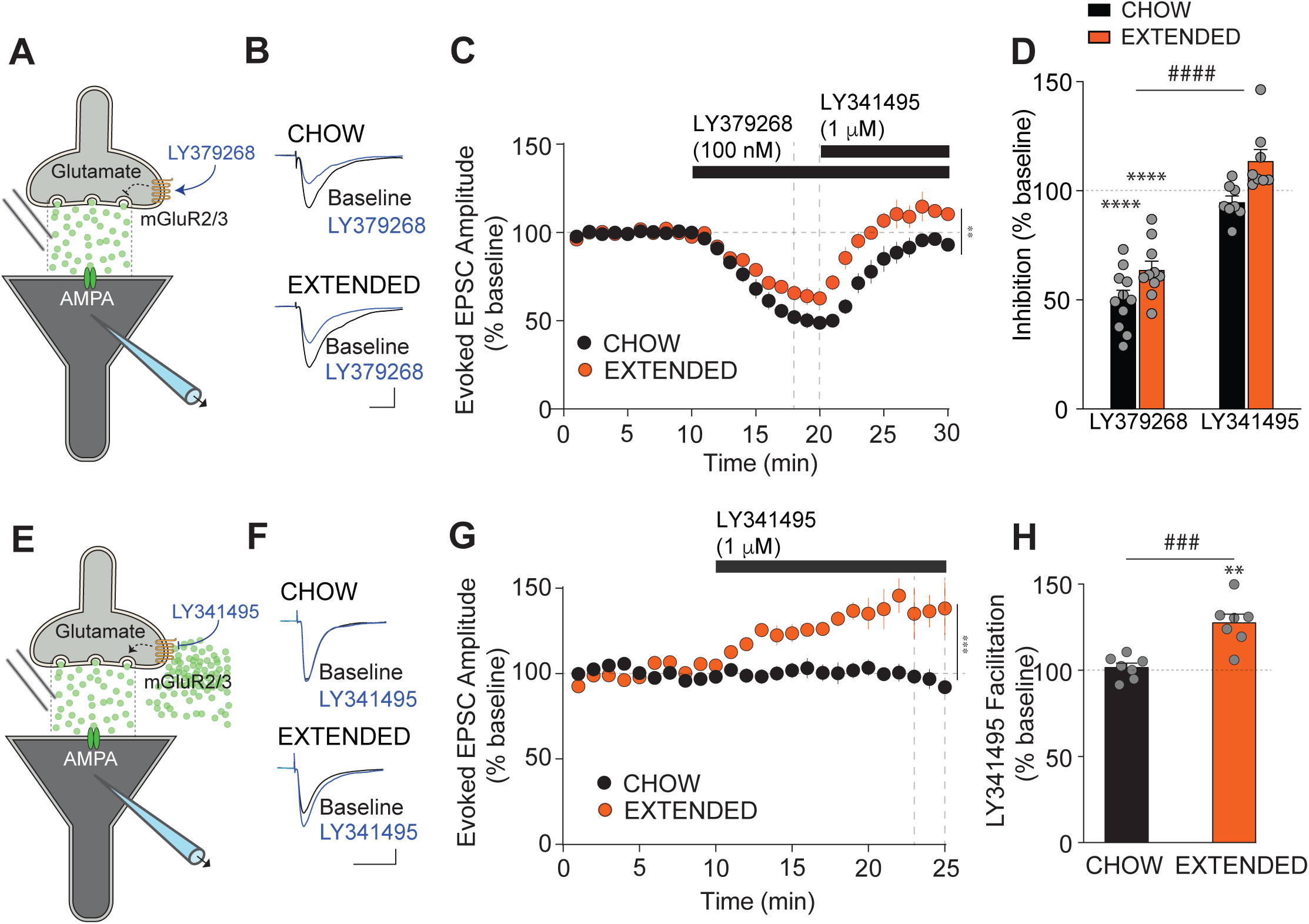
Glutamatergic synapses of lOFC pyramidal neurons from extended access rats are tonically suppressed by glutamate acting via extrasynaptic mGluR2/3 receptors. **(A)** Experimental schematic illustrating the effect of the mGluR2/3 agonist, LY379268 on evoked AMPA EPSCs. **(B)** Example traces of evoked AMPA EPSCs before (Baseline) and during application of LY379268 (100 nM) in lOFC neurons from chow or extended access rats. Scale bar: 500 pA, 10 ms. **(C)** Time plot of evoked EPSC amplitude before and during application of LY379268 (100 nM), and then after further addition of the mGluR2/3 receptor antagonist, LY341495 (1 μM) in lOFC neurons from chow (n = 11/4) or extended access rats (n = 10/4). Dotted vertical lines show the time period analyzed in (D). There was a significant Diet x Time interaction, F(29,494) = 4.097, P < 0.0001 and a main effect of diet, F(1,18) = 11.03, P = 0.0038**. **(D)** Bar graph quantifying the percentage inhibition of evoked EPSC amplitude by LY379268 (100 nM) and its reversal by LY341495 (1 μM) in Chow and Extended access rats. LY379268 produced significantly more inhibition of evoked EPSC amplitude in lOFC neurons from chow compared to extended access rats. A mixed-effects model (REML) indicated no diet x drug interaction (F(1,33) = 0.4314, P =0.5159), but main effects of diet (F(1,33) = 14.75, P= 0.0005^###^ and drug (F(1,33)=125.9, P < 0.0001^####^). There was a significant LY379268-induced inhibition of EPSCs compared to baseline in lOFC neurons from chow (n = 11/4, t_(10)_ =12.06, P <0.0001****) and extended access rats (n = 10/4, t_(9)_ = 8.997, P <0.0001****). **(E)** Experimental schematic illustrating an extrasynaptic glutamate tone acting via presynaptic mGluR2/3 receptors to suppress synaptic glutamate release. The effect of the mGluR2/3 antagonist, LY341495 was examined on evoked AMPA EPSCs to unmask this suppression. **(F)** Example traces of evoked AMPA EPSCs before (Baseline) and during application of LY341495 (1 μM) in lOFC neurons from chow and extended access rats. Scale bars: 250 pA, 10 ms. **(G)** Time course of evoked EPSC amplitude before and during application of the mGluR2/3 antagonist, LY341495 (1 μM) in lOFC neurons from chow (n = 7/4) and extended access rats (n = 7/3). A mixed-effects model (REML) indicates a significant diet x time interaction (F(24,296) = 12.40, P < 0.0001) and a significant diet effect: (F(1,13) = 24.35, P = 0.0003***). **(H)** Bar graph quantifying the percentage facilitation of evoked EPSC amplitude by LY341495 in lOFC neurons from chow (n = 7/4) and extended (n = 7/3) access rats. There was a significant difference in effect by LY341495 on EPSC amplitude between lOFC neurons from chow and extended access rats (t_(12)_ = 4.57, P = 0.0006^###^). There was a significant LY341495-induced facilitation of EPSCs over baseline in lOFC neurons from extended access rats (t_(6)_ = 5.424, P = 0.0016**), but not chow access rats (t_(6)_ = 0.6378, P = 0.5471). Bars represent mean ± s.e.m with individual values overlaid.

### An obesogenic diet induces hypertrophy of astrocytes and impairs their ability to transport glutamate

So far, we have demonstrated that during obesity, lOFC pyramidal neurons receive enhanced glutamate from an extrasynaptic source. There is an increasing body of work implicating astrocytes in the regulation of synaptic transmission (Dallérac et al., 2018; Hirase et al., 2014). Given that astrocytes account for the majority of glutamate transport in the brain and are altered in diet-induced obesity (García-Cáceres et al., 2013), we surmised that an obesogenic diet impairs astrocytes and their ability to uptake glutamate. We first examined immunofluorescence of the astrocyte marker, glial fibrillary acidic protein (GFAP) within the OFC of chow, restricted or extended access rats (Figure 5A). Compared to the OFC of chow or restricted access rats, we found an increase in the intensity and area of GFAP in extended access rats (Figure 5A,B,C,E,F). However, there was no change in the number of GFAP+ astrocytes (Figure 5D,G). To investigate this further, we examined the finer morphology of astrocytes using two-photon fluorescence microscopy (Figure 5H). OFC astrocytes were first labelled *in-vitro* with the astrocyte-specific dye, SR101 (20 μM, 20 min). Individual astrocytes were then patch-clamped under fluorescence and filled with the large fluorescent dye, FITC- dextran (3-5 kDa) (Figure 5H). In addition to observing typical astrocyte morphology, astrocytes were confirmed electrophysiologically, indicated by a lack of cell firing, low input resistance and passive membrane conductance (Figure S4). Astrocytes from the lOFC of extended access rats had significantly larger astrocyte territories (4407 ± 203 μm^2^) compared to those from chow access rats (3654± 214 μm^2^) (Figure 5H-J). However, there was no change in branching indicated by the Schoenen ramification index (Figure 5K). Together, these data indicate that extended access to an obesogenic diet induces hypertrophy of OFC astrocytes.

**Figure 5:**
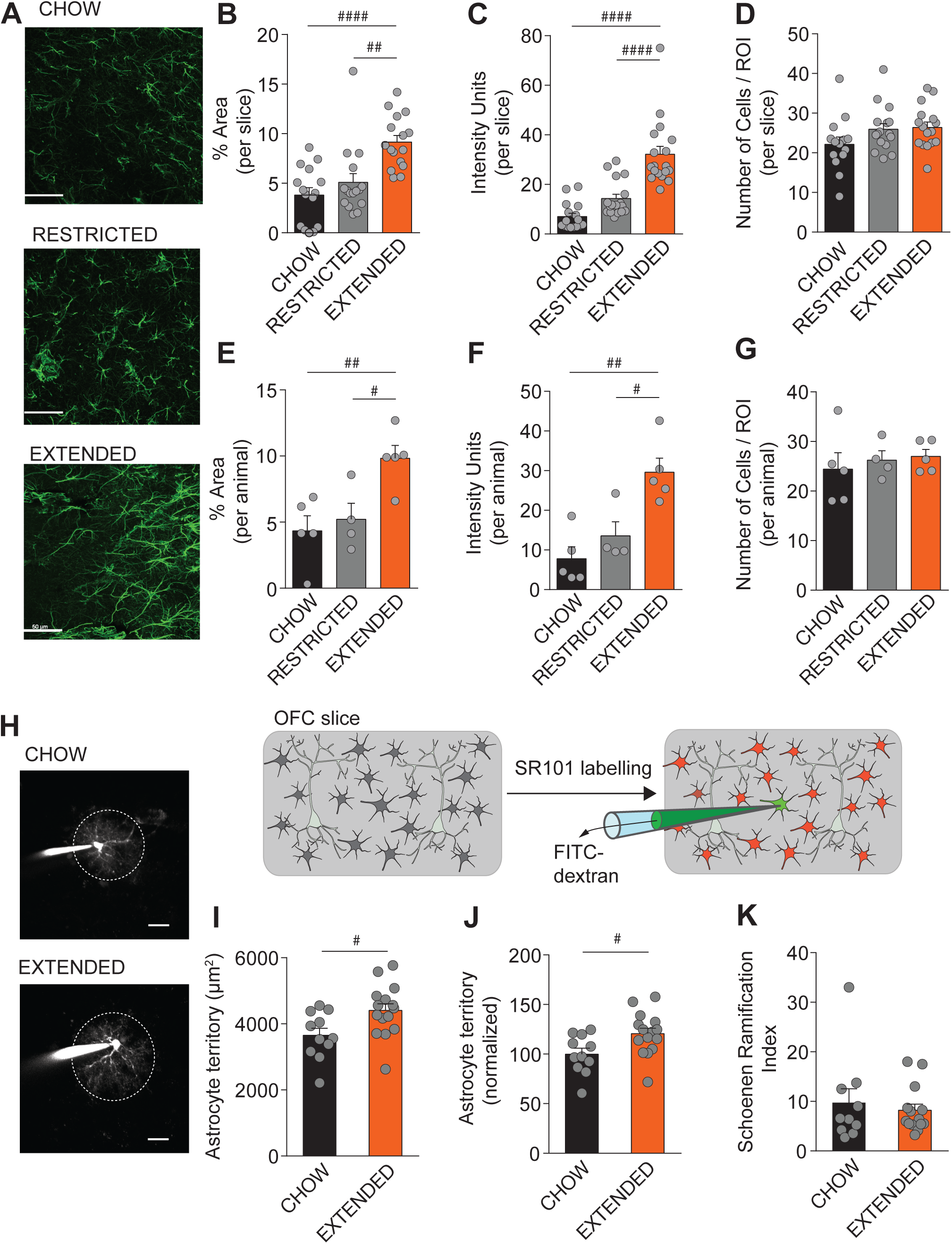
OFC astrocytes are hypertrophic following extended exposure to an obesogenic diet. **(A)** Representative images illustrating GFAP immunofluorescence in the lOFC of chow (upper), restricted (middle) and extended (lower) access rats. Scale bar, 50 µm. **(B)** Quantification of percentage area of fluorescence of GFAP from slices containing the lOFC of chow (n= 12 slices/5 rats), restricted (n = 16 slices/4 rats) or extended (n = 14/5 rats). GFAP staining in lOFC of extended access rats is greater than that of that from chow (P<0.0001^####^) or restricted access rats (P<0.01^##^), One way ANOVA: F(2,44) = 13.29, P < 0.0001). **(C)** Quantification of relative intensity units of GFAP fluorescence from slices containing the lOFC of chow (n= 16 slices/5 rats), restricted (n = 17 slices/4 rats) or extended (n = 18/5 rats). GFAP staining in lOFC of extended access rats is greater than that of that from chow (P<0.0001^####^) or restricted access rats (P<0.0001^####^) One way ANOVA: F(2,48) = 31.85, P < 0.0001). **(D)** Quantification of the number of GFAP expressing cells from slices containing the lOFC of chow (n= 12 slices/5 rats), restricted (n = 16 slices/4 rats) or extended (n = 14/5 rats). There is no significant difference in the number of GFAP+ cells in lOFC of chow, restricted or extended access rats (One way ANOVA: F(2,43) = 2.207, P= 0.1223). **(E)** Quantification of percentage area of fluorescence of lOFC GFAP from 3-4 slices averaged by animal. GFAP staining in lOFC of extended access rats (n = 5) is greater than that of that from chow (n = 5, P<0.01^##^) or restricted access rats (n = 4, P<0.05^#^) One way ANOVA: F(2,11) = 7.526, P 0.0087). **(F)** Quantification of relative intensity units of lOFC GFAP fluorescence from 3-4 slices averaged by animal. GFAP staining in lOFC of extended access rats (n = 5) is greater than that of that from chow (n = 5, P<0.01^##^) or restricted access rats (n = 4, P<0.05^#^), One way ANOVA: F(2,11) = 12.08, P = 0.0017). **(G)** Quantification of the number of GFAP expressing cells from 3-4 slices averaged by animal. There is no significant difference in the number of GFAP+ cells in lOFC of chow (n = 5), restricted (n = 4) or extended access rats (n = 5; One way ANOVA: F(2,11) = 0.314, P= 0.737). **(H)** Representative images illustrating fluorescent dye-filling of astrocytes within layer II/III of the lOFC in slices from chow and extended access rats. Scale bars: 10 µm. Inset, experimental schematic outlining imaging of astrocyte processes. Coronal slices containing OFC were first loaded with the astrocyte-specific dye, sulfurhodamine 101 (SR101, 20 μM, 20 min). SR101 labelled astrocytes were then patch-clamped under fluorescence, filled with the dye, FITC-dextran to label the fine processes and then imaged using 2-photon microscopy. **(I)** Area of individual astrocyte territory in lOFC from chow (n = 11/4) and extended (n = 15/4) access rats. Astrocyte territory is significantly increased in lOFC from extended access rats compared to chow access rats (t_(24)_ = 2.514, P = 0.019^#^). **(J)** Area of individual astrocyte territory normalized to average. Normalized astrocyte territory is significantly increased in lOFC from extended access rats (n = 15/4) compared to chow access rats (n = 11/4) (t_(24)_ = 2.514, P = 0.019^#^). **(K)** Astrocyte process arborization indicated by the Schoenen ramification index. Astrocyte ramification is not different between chow (n = 10/4) and extended (n = 14/4) access rats (t_(22)_ = 0.532, P = 0.6003). Two cells were omitted due to high background fluorescence. Bars represent mean ± s.e.m with individual values overlaid.

We next explored whether hypertrophic astrocytes in the OFC were associated with impaired glutamate transporter 1 (GLT-1) function, an astrocytic transporter responsible for approximately 90% of glutamate reuptake in the cortex (Armbruster et al., 2016; Danbolt et al., 1992; Hanson et al., 2016; Voutsinos-Porche et al., 2003). Patch-clamping from SR101-labelled lOFC astrocytes in artificial cerebrospinal fluid (ACSF) containing caged Rubi-Glutamate (30 μM), we evoked the focal action of glutamate via flash photolysis (473 nm LED, 2 ms, Figure 6A). An astrocytic glutamate transporter current was isolated in the presence of synaptic blockers and by subtracting the residual current following application of TBOA (30 μM) (Figure 6B). Glutamate transporter currents in lOFC astrocytes from extended access rats were significantly reduced compared to those from chow access rats (Figure 6B,C). In contrast to the lOFC, we found that the magnitude of astrocytic GLT-1 currents was not different between chow and extended access rats in the prelimbic and motor cortex (Figure S1E,F). We further examined GLT-1 protein with western blots, but observed no difference in GLT-1 levels between the lOFC of chow or extended access rats (Figure 6D,E). Together, these results indicate impairment in the function but not quantity of GLT-1 in lOFC astrocytes from obese rats.

**Figure 6:**
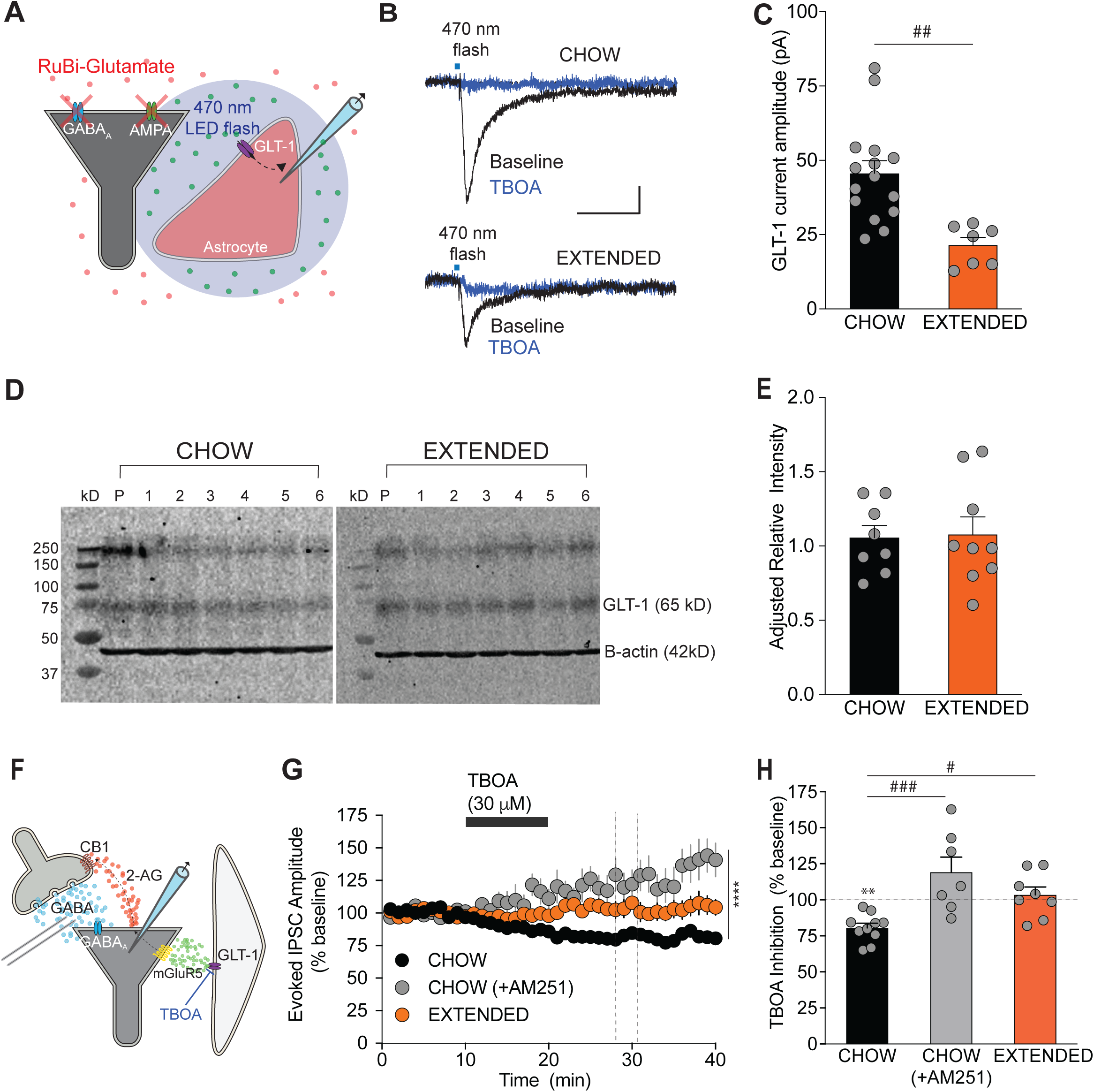
OFC astrocytes have impaired glutamate transport following extended exposure to an obesogenic diet. **(A)** Experimental schematic illustrating the recording of glutamate transporter currents in astrocytes. SR101-labelled astrocytes were patched-clamped at –80 mV with caged Rubi-Glutamate (30 μM) in the extracellular ACSF. Glutamate was focally activated on the astrocyte via a blue LED (470 nm) light flash. A glutamate transporter current was isolated in the presence of synaptic blockers. (**B**) Example traces showing astrocyte glutamate transporter currents evoked via flash photolysis (470 nm) of RuBi-Glutamate (30 μM) before and during application of the glutamate transport blocker, TBOA (30 μM) in the lOFC of chow and extended access rats. Scale bar: 10 pA, 25 ms. (**C**) Bar graph quantifying the amplitude of flash-evoked GLT-1 currents in astrocytes. GLT-1 current amplitude is significantly reduced in lOFC astrocytes from extended (n = 7/4) compared to chow (n = 15/5) access rats (t_(20)_ = 3.664, P = 0.0015^##^). (**D**) Example immunoblots of GLT-1 (65 kD) from the OFC of chow or extended access rats. Beta-actin (42 kD) is the loading control protein. Numbers above lanes refer to OFC samples from different chow or extended access rats. Pooled control (P) contains mixed brain tissue from cortex, hippocampus and striatum serving as a quality control. (**E**) Western blot analyses of total GLT-1 protein levels in the OFC of chow (N = 8) or extended (N = 9) access rats. There is no difference in GLT-1 presence between diet conditions (t_(15)_ = 0.159, P = 0.88). (**F**) Experimental schematic illustrating a glutamate transporter-induced inhibitory LTD. Blockade of glutamate transport with TBOA (30 μM) leads to an excess of glutamate in the extrasynaptic space, which indirectly drives retrograde endocannabinoid release onto GABA synapses. (**G**) The glutamate transport inhibitor, TBOA (30 μM) induces a long-term depression of evoked IPSCs onto lOFC pyramidal neurons from chow access rats (n =10/8). This effect is inhibited when slices are pre-treated with the CB1 receptor antagonist, AM251 (3 μM, n = 7/5). Furthermore, TBOA-induced LTD is occluded in lOFC neurons from extended access rats (n = 8/3). Mixed effects model (REML): F(1.226,63.14) = 102.9, P < 0.0001). Dotted vertical lines indicate the time period analyzed in (H). (**H**) Bar graph quantifying the percentage change of evoked IPSC amplitude during TBOA application in slices from untreated (n = 10/8) and AM251-treated (3 µM, n = 7/5) neurons from chow or extended access rats (n/N = 8/3). TBOA produces a long-term suppression of evoked IPSC amplitude in lOFC neurons from chow access rats, which is blocked in the presence of AM251 and absent in extended access rats. One way ANOVA: F(2,22) = 9.804, P = 0.0009, Tukey’s multiple comparison test: chow vs. chow + AM251, P<0.001^###^, chow vs. extended, P<0.05^#^. Compared to baseline, TBOA significantly decreased evoked IPSC amplitude in lOFC neurons from chow access rats (t(9) = 6.307, P=0.0001***). However, evoked IPSC amplitude was unchanged in lOFC neurons from chow access animals pre-treated with AM251 (t(6) = 1.824, P = 0.1180), or from extended access rats (t(7) = 0.6321, P = 0.5474). Cells that did not reach the 27-31 min analysis period following TBOA application were excluded. Bars represent mean ± s.e.m with individual values overlaid.

We next tested the hypothesis that impairment of astrocytic glutamate transport leads to an endocannabinoid-mediated LTD of GABAergic transmission (Figure 6F). Similar to TBS- induced LTD and mGluR-LTD, application of TBOA (30 μM, 10 min) suppressed evoked IPSC amplitude in lOFC neurons from chow access rats, which persisted for at least 20 min after washout of the drug (Figure 6G,H). This long-lasting suppression of inhibition was blocked by pre-treatment with AM251 (3 μM), indicating that this glutamate transporter-induced effect is endocannabinoid-mediated. The TBOA-induced suppression of inhibition was absent in lOFC neurons from extended access rats, consistent with an occlusion by extrasynaptic glutamate resulting from impaired transporter function (Figure 6G,H). Taken together, these results indicate that inhibition of astrocytic glutamate transport can influence lOFC inhibitory synapses via an endocannabinoid-mediated LTD, and this effect is impaired during diet-induced obesity.

### Restoration of glutamate homeostasis with N-acetylcysteine reverses the cascade of obesity-induced synaptic deficits

Because altered GABAergic synaptic transmission in lOFC pyramidal neurons from extended access rats is mediated upstream by impaired glutamate transport, we surmised that restoration of glutamate homeostasis *in vitro* could reverse the cascade of synaptic changes observed with diet-induced obesity. We tested the effect of N-acetylcysteine (NAC), a cystine prodrug reported to enhance GLT-1 activity and restore extrasynaptic glutamate concentration (Kupchik et al., 2012; Reissner et al., 2015). Following a 90 min pre-treatment of slices with NAC (500 μM), glutamate transporter currents were significantly enhanced in astrocytes of extended access rats, restoring their magnitude to that observed in chow access rats (Figure 7A). This was associated with a reduction in high frequency stimulation-evoked NMDA EPSC decay time and a restoration in the ability of TBOA to enhance this measure (Figure 7B). Taken together, NAC enhances the function of astrocytic glutamate transporters, leading to a reduction in extrasynaptic glutamate onto lOFC pyramidal neurons of obese rats.

**Figure 7:**
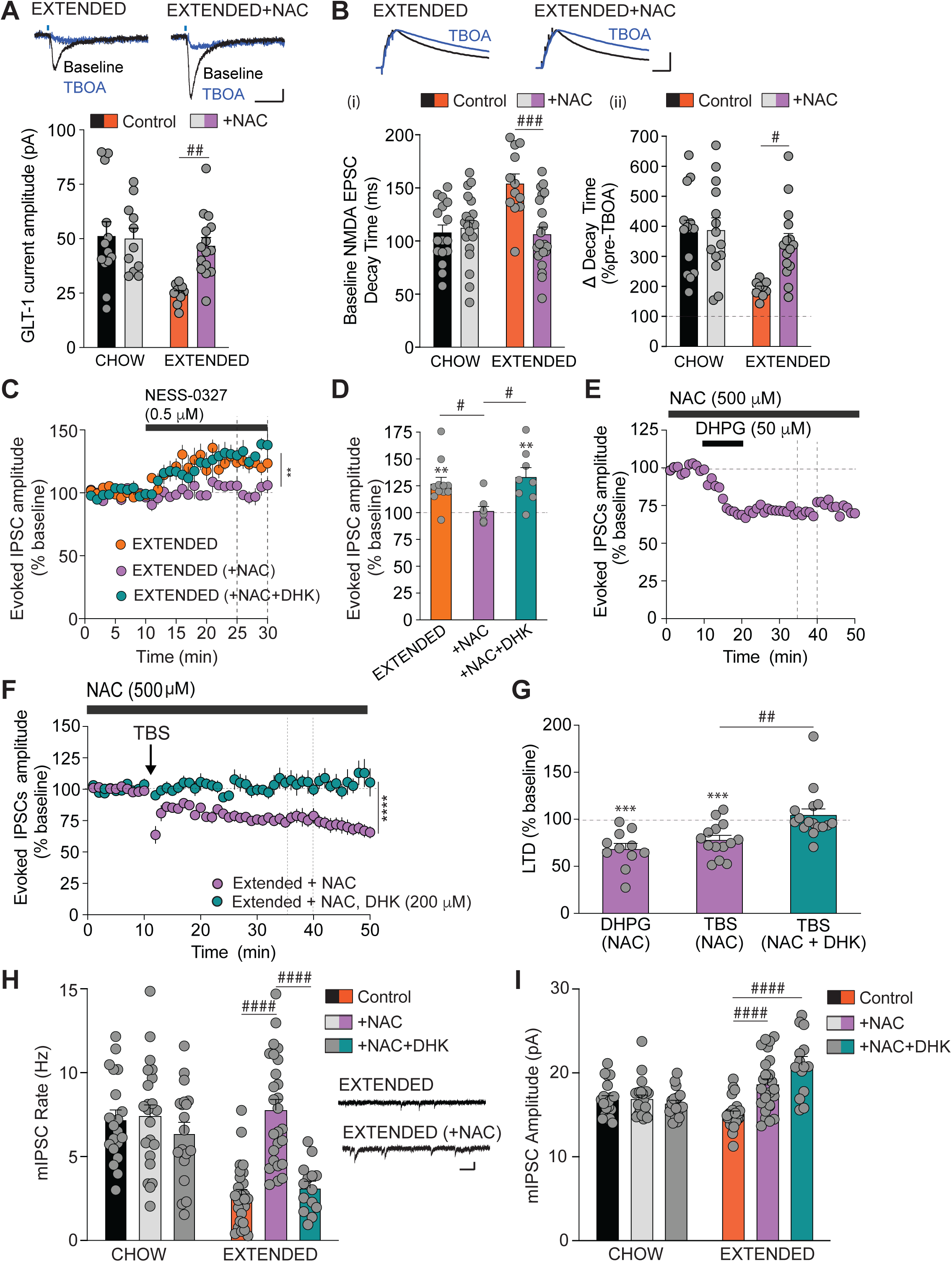
Restoration of glutamate homeostasis via astrocytic GLT-1 reverses the cascade of diet-induced alterations impairing inhibitory synaptic plasticity. (**A**) NAC treatment enhances GLT-1 current amplitude in astrocytes from extended access rats (Chow: n = 13/4, Chow + NAC: n = 11/4, Extended: n = 10/5, Extended + NAC: n = 15/5). There was a significant diet x drug interaction (F(1,45) = 5.847, P=0.197), with main effects of diet (F(1,45) = 4.698, P = 0.0355) and drug (F(1,45) = 9.675, P = 0.0032). A Sidak’s multiple comparison’s test revealed a significant difference between control and NAC in the extended access group (P = 0.0044^##^). Inset: Example traces of GLT-1 currents from lOFC astrocytes in untreated and NAC-treated slices from extended access rats. Scale bars: 10pA, 25 ms. (**B**) (i) NAC treatment reduces baseline NMDA EPSC half-decay time in lOFC pyramidal neurons from extended access rats comparable to chow rats (Chow: n = 16/6, Chow + NAC: n = 20/6; Extended: n = 12/4, Extended + NAC: n = 22/7; Interaction: F(1,66) = 11.66, P = 0.0011, diet effect (F(1,66) = 6.851, P= 0.0110, Drug effect: F(1,66) = 0.8,043, P = 0.0061). A Sidak’s post-hoc test indicated a significant difference between control and NAC in the extended access group (P<0.001^###^). (ii) Percentage change in decay time of evoked NMDA EPSCs in response to application of the glutamate transporter inhibitor, TBOA (30 µM, 10 mins) in lOFC neurons from chow (n = 13/4), extended access (n =9/3), and NAC treated slices from chow (n = 14/4) and extended access rats (n = 15/6). There was a significant interaction of diet and NAC on the change in decay time by TBOA (F(1,47) = 4.055, P = 0.49) and a main effect of diet (F(1,47) = 10.01, P = 0026) and a main effect of NAC (F(1,47) = 4.811, P = 0.033). A Sidak’s post-hoc test reveals a significant difference between control and NAC in the extended access group (P = 0.0129^#^). Inset: Example traces of evoked NMDA EPSCs normalized to peak amplitude from lOFC pyramidal neurons in response to 5X100 Hz stimulation in untreated and NAC-treated slices from rats with extended access to an obesogenic diet. Artefacts minimized for clarity. Scale bars: 0.5, 50 ms. (**C**) NAC treatment (n = 8/6) abolishes the facilitation of evoked IPSC amplitude unmasked by the neutral CB1 receptor antagonist, NESS-0327 (0.5 μM) in lOFC slices from extended access rats (n = 11/9). The effect of NAC is reversed by the selective GLT-1 blocker, dihydrokainic acid, (DHK, 200 μM, n = 8/6). A mixed-effects model (REML) indicates a significant NAC treatment x time interaction (F(58,686) = 2.441, p < 0.0001^####^) and a main effect of drug treatment (F(2,24) = 6.593, P < 0.001^##^). Broken vertical lines indicate time period analyzed in (D). (**D**) Bar graph quantifying evoked IPSC amplitude during NESS-0327 (0.5 μM) application in untreated vs. NAC treated slices from extended access rats. There was a significant increase of IPSC amplitude over baseline with NESS-0327 treatment in lOFC neurons of extended access rats (n=11/9; t_(10)_=4.235, P=0.0017**), but not when slices were treated with NAC (n=8/6, t_(7)_=0.338, P=0.745). The effect of NAC was prevented by DHK (n = 8/6, t_(7)_=3.755, P=0.0071**). A one-way ANOVA indicates a significant group difference (F(2,24) = 5.646, p = 0.0098) and a Tukey’s post-hoc test indicates significant differences between control and NAC (P<0.05^#^) and NAC and NAC+DHK (P<0.05^#^) in extended access rats. (**E**) Time plot of evoked IPSC amplitude before and after application of the mGluR1/5 agonist, DHPG in lOFC neurons from extended access rats pre-treated with NAC (n =11/8). NAC treatment restores the ability to induce mGluR-LTD. Broken vertical lines indicate time period analyzed in (G). (**F**) Time plot of evoked IPSC amplitude before and after TBS in lOFC neurons from extended access rats pre-treated with NAC alone (n = 14/7), or NAC with the selective GLT-1 inhibitor, DHK (200 μM) (n = 16/12). TBS-LTD was restored in lOFC neurons from extended access rats pre-treated with NAC, but not in the additional presence of DHK (RM REML: Time x drug interaction: F(48, 1270) = 5.302, P <0.0001; drug effect: F(1, 28) = 20.96, P < 0.0001****)). Dotted vertical lines indicate the 25-30 min time period following TBS analyzed in (G). (**G**) Bar graph quantifying evoked IPSC amplitude following application of DHPG (n = 16/12) or TBS (n= 14/7) in NAC treated slices from extended access rats. NAC treatment restores the ability to induce TBS-LTD (t_(13)_=4.564, P=0.0005***, one-sample t-test difference from baseline) or mGluR-LTD (t_(10)_=5.303, P=0.0003***, one-sample t-test difference from baseline) in lOFC neurons from extended access rats. The rescue of TBS LTD was prevented in the presence of DHK (t_(15)_=0.673, P=0.5024, one-sample t-test compared to baseline). There was a significant difference between groups (One way ANOVA F(2,38) = 9.992, P = 0.003, Tukey’s post-hoc test: NAC-TBS vs NAC-TBS+DHK, P<0.01^##^). (**H**) Effect of NAC on miniature IPSC rate in lOFC pyramidal neurons from chow or extended access rats (2-way ANOVA: diet x drug interaction: F(2,122) = 10.65, P<0.0001), diet effect: F(1,122 = 26.88, P <0.0001, drug effect: F(2,122) = 16.03, P<0.0001). A Tukey’s post-hoc test indicates that NAC treatment enhances mIPSC rate in lOFC neurons from extended access rats (extended (n = 26/3) vs. extended+NAC (n = 26/3), p<0.0001^####^), but had no effect in chow access rats (chow (n =20/3) vs. chow+NAC (n=24/3), P>0.05). Further, mIPSC rate was significantly different between lOFC neurons of chow and extended access rats (P <0.0001^####^). Furthermore, controls were not significantly different from DHK + NAC treatment in either chow (n = 16/3; P>0.05) or extended access rats (n = 14/3; P>0.05). Inset: Example traces of mIPSCs in untreated and NAC-treated slices from extended access rats. Scale bar: 20pA, 100 ms. (**I**) Effect of NAC on miniature IPSC amplitude in lOFC pyramidal neurons from chow or extended access rats (2-way ANOVA: diet x drug interaction: F(2,122) = 16.22, P<0.0001), diet effect: F(1,122 = 12.26, P = 0.0006, drug effect: F(2,122) = 12.51, P<0.0001). A Tukey’s post-hoc test indicates that NAC did not alter mIPSC amplitude in lOFC neurons from chow access rats (p>0.05), but enhanced this measure in lOFC neurons from extended access rats (P<0.0001^####^). Furthermore, while there was no significant difference between chow control and DHK +NAC treatment (P>0.05), in extended access rats, DHK +NAC treatment significantly increased mIPSC amplitude compared to controls (P<0.0001^####^), but was not significantly different from NAC treated (P>0.05). Bars represent mean ± s.e.m with individual values overlaid.

We then tested whether NAC could abolish the mGluR5-induced endocannabinoid tone and restore LTD at inhibitory synapses. Pre-treatment with NAC prevented the facilitation of evoked IPSC amplitude induced by NESS-0327 (Figure 7C,D) and restored both mGluR-LTD and TBS-LTD at inhibitory synapses onto lOFC neurons (Figure 7E,F,G). To confirm that NAC restores endocannabinoid tone and TBS-LTD through manipulation of glutamate transport, we re-examined the effect of NAC in the combined presence of the selective GLT-1 blocker, dihydrokainic acid (DHK). DHK (200 μM) prevented NAC-induced restoration of endocannabinoid tone (Figure 7C,D) and blocked TBS-induced LTD (Figure 7F,G), indicating that NAC rescues these synaptic changes by restoring astrocyte glutamate clearance. Finally, pre-treatment with NAC significantly enhanced the frequency of mIPSCs onto lOFC neurons from extended access rats, similar to that observed in chow access rats (Figure 7H). However, NAC had no influence on mIPSC frequency in chow access rats, suggesting an action specific to the obese state. Additionally, application of the GLT-1 inhibitor prevented the NAC-induced restoration of mIPSC frequency (Figure 7H), further confirming a mechanism of action via GLT-1. Together, these data indicate that by restoring glutamate transporter function and extrasynaptic glutamate levels, NAC abolishes endocannabinoid tone and the occlusion of endocannabinoid-mediated LTD, thereby rescuing the impairment of GABAergic synaptic transmission in obese rats.

## Discussion

These observations provide a number of new insights that expand our understanding of how obesogenic diets alter endocannabinoid function in the brain. Of primary importance, our data identify a novel astrocyte-synaptic mechanism in the lOFC that becomes disrupted in obesity. Following long-term exposure to an obesogenic diet, lOFC astrocytes undergo hypertrophy and this is associated with impairment of their predominant glutamate transporter, GLT-1. This leads to enhanced extrasynaptic glutamate, which generates an endocannabinoid tone via mGluR5 activation to induce a LTD of GABA synaptic transmission onto lOFC pyramidal neurons. Importantly, this cascade of synaptic deficits could be reversed by restoring glutamate homeostasis with the nutritional supplement, NAC. The data presented here demonstrate that an obesogenic diet acts via a neuron-astrocyte interaction to induce heterosynaptic plasticity of excitatory and inhibitory synapses within the lOFC. Furthermore, these effects are not ubiquitous in the cortex as we did not observe these synaptic changes in the primary motor cortex or mPFC.

### An obesogenic diet alters astrocyte structure and function in the lOFC

Extended access to an obesogenic diet led to structural hypertrophy of OFC astrocytes, whereby the area of their territory expanded by ∼ 20%. This was accompanied by increased GFAP immunoreactivity, which may be indicative of reactive astrocytes, although not an absolute marker (Escartin et al., 2021). This is consistent with astrocyte reactivity reported in the hypothalamus following short-term obesogenic diet exposure (Thaler et al., 2012), as well as after diet-induced obesity is established (Thaler et al., 2012; Zhang et al., 2017). Hypothalamic astrocytes are reported to undergo shortening of their processes as measured with GFAP immunostaining (Zhang et al., 2017). However, the large GFAP+ primary branches only account for 10-20% of the total volume of astrocyte arborisation (Bushong et al., 2002). Therefore, to more thoroughly examine the fine processes of astrocytes not labelled by GFAP, we performed fluorescent dye-filling of OFC astrocytes using two-photon microscopy. We observed lengthening of astrocytic processes without an increase in number of GFAP+ cells. Given that the ability of astrocytes to impact synaptic function relies on their physical interaction with synapses (Pannasch et al., 2014; Papouin et al., 2017), changes in astrocyte territory are likely to modify basal synaptic activity. Indeed, we observed altered kinetics of extrasynaptic glutamate action in the lOFC of obese rats. Enhanced extrasynaptic glutamate was likely due to impaired clearance via reduced astrocytic GLT-1 function. Consistent with this, astrocytic glutamate transporter currents were significantly reduced in the lOFC of obese rats. These currents were abolished in the presence of the non-selective glutamate transport blocker, DL-TBOA, indicating that the astrocytic glutamate transporters, GLAST and/or GLT-1 were involved. Given the respective EC_50_ of DL-TBOA for these transporters is 70 μM and 6 μM (Shimamoto et al., 2000), the 30 μM concentration of DL-TBOA used in this study should have predominantly targeted GLT-1. In addition to reduced GLT-1 function, we observed an increase in length of astrocytic processes with no change in quantity of GLT-1 protein, suggesting that GLT-1 may be diluted along the processes of lOFC astrocytes of obese rats. Other studies using obesogenic diets have reported mixed findings in GLT-1 number, demonstrating either a reduction (Linehan et al., 2018; Tsai et al., 2018) or upregulation of GLT-1 in hypothalamic or hippocampal tissue (Cano et al., 2014; Valladolid-Acebes et al., 2012). This discrepancy may be associated with the density of astrocytic GLT-1 transporter in reference to altered astrocytic morphology (Zhang et al., 2017). Astrocyte morphology can influence glutamate transport (Oliet et al., 2001; Pannasch et al., 2014), and GLT-1 can rapidly diffuse within the membrane to influence EPSC kinetics (Murphy-Royal et al., 2015). Thus, it is conceivable that a diluted presence of GLT-1 on extended astrocytic processes in the OFC of obese rats may have a negative impact on synaptic glutamate homeostasis by disconnecting glutamate reuptake from synaptic release.

A consequence of reduced astrocytic glutamate transport in the OFC of obese rats was the tonic presence of glutamate in the extrasynaptic space. This excess of extrasynaptic glutamate indirectly led to downstream suppression of synaptic GABA (via mGluR5-induced endocannabinoid signalling) and glutamate (via mGluR2/3-mediated autoinhibition). Thus, by altering extrasynaptic glutamate transport, an obesogenic diet engages distinct mGluR mechanisms to modulate inhibitory and excitatory synaptic transmission. It should be noted that while extrasynaptic transmission by glutamate can directly influence synaptic release, it may also act via non-synaptic processes to alter neuronal function. In particular, glutamate can act on neighbouring astrocytes to modulate gliotransmitter release, which participates in shaping the activity and plasticity of neurons (Dallérac et al., 2018).

### An obesogenic diet disrupts endocannabinoid-mediated inhibitory synaptic plasticity in the lOFC

Rats with extended access to a cafeteria diet exhibited decreased GABAergic synaptic transmission onto layer II/III pyramidal neurons within the lOFC, consistent with our previous findings (Thompson et al., 2017). This was likely due to a suppression in presynaptic GABA release, as we observed a change in the rate, but not amplitude of mIPSCs. Furthermore, this presynaptic inhibition was associated with the presence of endocannabinoid tone, as a CB1 receptor antagonist blocked the obesity-induced reduction of mIPSC rate and unmasked a facilitation of evoked IPSCs. These synaptic changes were associated with an inability to induce endocannabinoid-mediated LTD, requiring activation of mGluR5 by extrasynaptic glutamate, suggesting that a prior experience of such plasticity mediates the suppression of inhibition by endocannabinoids. Multiple lines of evidence support this notion. An mGluR5 antagonist blocked endocannabinoid-mediated TBS-induced LTD in naive rats. Moreover, application of an mGluR1/5 agonist induced an LTD in chow-fed, but not obese rats. Finally, in obese rats, application of the mGluR5 antagonist blocked endocannabinoid tone unmasked by the CB1 receptor antagonist and mimicked this tone when applied alone. These data suggest that an obesity-induced endocannabinoid tone is the result of upstream activation of mGluR5, rather than a downstream alteration of endocannabinoid synthetic or degradative enzymes.

Obesity-induced impairment in astrocytic glutamate reuptake indirectly led to a reduction of GABAergic synaptic transmission in the lOFC. To confirm that GLT-1 function is directly linked to an extrasynaptic glutamate-driven and endocannabinoid-mediated suppression of GABAergic transmission, we showed that blockade of GLT-1 induced an LTD of GABAergic transmission in the lOFC of chow access rats. This was endocannabinoid-mediated as it was prevented in the presence of a CB1 receptor antagonist. Importantly, this LTD induced by glutamate transport blockade was absent in lOFC neurons of obese rats, suggesting an occlusion by an existing extrasynaptic glutamate tone. Therefore, our results suggest that a failure in glutamate reuptake by astrocytes can heterosynaptically modulate GABAergic transmission via recruitment of endocannabinoids. While synaptic glutamate spillover can induce endocannabinoid-mediated suppression of GABAergic synapses in neurons under physiological conditions (Chevaleyre and Castillo, 2004; Drew et al., 2008), to our knowledge, the present study is the first to demonstrate that astrocyte dysfunction can drive this phenomenon during a pathophysiological state such as obesity.

There is considerable evidence linking diet to alterations in brain endocannabinoid function (Gómez-Pinilla, 2008). Importantly, endocannabinoids are synthesized from essential fatty acids which are strictly derived from dietary sources (Simopoulos, 2016). A diet deficient in omega-3 fatty acids impairs synaptic plasticity within a number of brain regions, and this is associated with behavioural impairments (Lafourcade et al., 2011; Manduca et al., 2017; Thomazeau et al., 2017). The cafeteria diet employed here was composed of approximately 50% fat from mostly saturated sources. While omega-3 fatty acids were present in chow given to all diet groups, rats with restricted or extended access to the cafeteria consume significantly less chow (Thompson et al., 2017). Given that the synaptic impairments observed herein only occurred in rats with extended access to the diet, it is more likely that the obese state rather than the diet *per se* contributed to the synaptic changes within the lOFC.

There are multiple consequences of elevated endocannabinoid tone in the lOFC of obese mice. Dendritic spine density is reduced following chronic cannabinoid exposure (Chen et al., 2013; Rubino et al., 2009). Notably, we previously observed a reduction of spine density on lOFC pyramidal neurons in rats given extended access to a cafeteria diet (Thompson et al., 2017). Thus, the persistent endocannabinoid tone induced by the diet may influence morphological features of lOFC pyramidal neurons. Elevated endocannabinoid tone may also disrupt the interplay of synaptic excitation and inhibition influencing cortical processing. We demonstrated that TBS-induced LTD at inhibitory synapses in the lOFC requires endocannabinoids through a canonical mGluR1/5-mediated mechanism in chow fed or naive rats. However, after diet-induced obesity, both mGluR1/5- and TBS-induced LTD were occluded. Suppressed inhibitory plasticity may disrupt the feedback and feed-forward circuits underlying the regulation of cortical network activity. Notably, deletion of CB1 receptors from OFC output neurons disrupts endocannabinoid regulation of terminals in the dorsal striatum, leading to habit formation (Gremel et al., 2016). Further, activation of CB1 receptors in the OFC influences cost-benefit decision making (Khani et al., 2015). Thus, disrupted endocannabinoid regulation of OFC principal output neurons may be important in driving inflexible behaviour.

It should be noted that the obesity-induced synaptic alterations observed here are not ubiquitous in the cortex. In addition to the OFC, we examined upstream GLT-1 function and downstream GABA transmission in two adjacent regions in the coronal plane, the M1 motor cortex and prelimbic cortex. By contrast, both these measures were unaltered between chow and extended access rats, suggesting specificity to the OFC. However, we cannot rule out similar changes in other regions of the cortex or brain.

### N-acetylcysteine restores the diet-induced alterations in excitatory and inhibitory plasticity

There is an increasing body of work indicating that NAC has efficacy in the treatment of neuropsychiatric conditions, including substance use disorder and compulsive behaviours (Baker et al., 2003; Brown et al., 2013; Grant et al., 2009, 2016; Moussawi et al., 2009). Furthermore, an accumulation of experimental evidence supports the therapeutic benefits of NAC treatment in obesity through its anti-inflammatory and antioxidant properties, as well as its ability to interfere with adipogenesis (Dludla et al., 2019). NAC can increase cystine-glutamate exchange and enhance glutathione synthesis, a potent antioxidant (McBean, 2002). In addition to activation of the cystine-glutamate exchanger, NAC appears to restore aberrant glutamate homeostasis associated with cocaine self-administration by enhancing glutamate transport via GLT-1 (Kupchik et al., 2012; Reissner et al., 2015). Here we showed that NAC could reverse all of the obesity-induced synaptic changes within the OFC by restoring GLT-1 activity. NAC directly enhanced the function of GLT-1 in astrocytes, which restored extrasynaptic glutamate transmission. This was associated with the elimination of endocannabinoid tone and a restoration of both mGluR- and endocannabinoid-mediated TBS-LTD, which ultimately restored GABA transmission. These restorative effects of NAC were blocked in the presence of the specific GLT-1 inhibitor, confirming its mechanism of action via GLT-1. However, we cannot rule out other potential mechanisms such as glutathione restoration, given that previous evidence indicates that GLT-1 function may be impaired under conditions of oxidative stress (Lauderback et al., 2001). Taken together, our data indicates that NAC can reverse the cellular alterations within the lOFC induced by long-term exposure to an obesogenic diet, suggesting possible therapeutic potential in reversing the synaptic deficits associated with obesity. While prior human and animal work has demonstrated efficacy of NAC in reducing food-intake and weight gain (Dludla et al., 2019; Hurley et al., 2016), whether it produces this effect via action within the OFC remains to be determined. Our findings have identified an extended cellular mechanism of action by NAC, which in addition to restoring glutamate homeostasis, may also normalize GABA levels within the brain.

In conclusion, our findings provide a novel cellular mechanism by which an obesogenic diet alters inhibitory and excitatory transmission within the OFC. By altering astrocytes and their ability to transport glutamate, an obesogenic diet disrupts glutamate homeostasis, resulting in excess glutamate in the extrasynaptic space. This indirectly leads to an endocannabinoid-mediated long-term suppression of GABAergic transmission. Together, this disinhibition may ultimately disrupt the excitatory-inhibitory balance in the OFC and resultant output from primary projection neurons of the OFC. This disruption in synaptic function may underlie behaviours associated with overeating, leading to a persistence of obesity. Importantly, these alterations in synaptic transmission are reversed by restoring glutamate homeostasis with NAC, highlighting the potential of this drug in obesity.

**Table 1.** Nutrient composition of Cafeteria diet (% Total diet)

| Food | Fat | Protein | Carbohydrate | Kcal/g |
| --- | --- | --- | --- | --- |
| Kraft Peanut butter | 53 | 20 | 27 | 6 |
| Kellog's Froot Loops | 3.7 | 3.7 | 89 | 4.1 |
| Doritos | 26 | 6 | 64 | 5.2 |
| Kirkland Hot Dogs | 25 | 12 | 3.5 | 3 |
| Tim Hortons Timbits | 16 | 5.3 | 53 | 3.7 |
| Lab Diet (5P14) Rat Chow | 4.5 | 24 | 49 | 3 |

## Funding and disclosure

This research was supported by a Koopmans Research Award, Canadian Institutes of Health Research operating grant (CIHR, FDN-147473) and a Canada Research Chair Tier 1 (950-232211) to SLB. Benjamin K Lau was supported by postdoctoral awards from the Cumming School of Medicine (Eyes High) and Alberta Innovates Health Solutions. The authors declare no competing financial interests.

## Acknowledgements

The authors would like to acknowledge the Hotchkiss Brain Institute optogenetic core facility and the advanced microscopy facility for their technical support.

## Data Accessibility

Data will be made available upon request.

## Author contributions

B.K.L. designed, performed and analyzed all electrophysiological experiments with supervision of S.L.B. C.M-R. performed astrocyte labeling experiments with supervision of G.G. and J.S.B. M.K. and M.Q. performed immunohistochemistry experiments and western blots under supervision of S.L.B. B.L.K. and S.L.B. wrote the manuscript.

## Figure Legends

**FIGURE S1. GABAergic transmission onto pyramidal neurons or GLT-1 currents in astrocytes are unaltered by an obesogenic diet in prelimbic or primary motor cortices.**

*(**A**)* Experimental schematic illustrating the the recording sites for (i) prelimbic or (ii) primary motor cortex relative to the lOFC.

*(**B**)* Example traces of mIPSCs onto (i) prelimbic cortex or (ii) primary motor cortex pyramidal neurons from chow or extended access rats. Scale bars: 50 pA, 200 ms.

*(**C**)* Bar graph quantifying the rate of mIPSCs onto (i) prelimbic or (ii) primary motor cortex pyramidal neurons from chow or extended access rats. There was no significant difference in mIPSC rate between diet groups for mIPSCs in the prelimbic cortex (t(24) = 0.79, P = 0.44; chow: n = 14/4, extended: n = 12/4) or the motor cortex (t(25) = 0.44, P = 0.66; chow: n = 15/5, extended: n = 12/4).

*(**D**)* Bar graph quantifying the amplitude of mIPSCs amplitude onto (i) prelimbic or (ii) primary motor cortex pyramidal neurons from chow or extended access rats. There was no significant difference in mIPSC amplitude between diet groups for mIPSCs in the prelimbic cortex (t(24) = 1.56, P = 0.13; chow: n = 14/4, extended: n = 12/4) or the motor cortex (t(25) = 0.52, P = 0.61; chow: n = 15/5, extended: n = 12/4).

*(**E**)* Example traces showing astrocyte glutamate transporter currents evoked via flash photolysis (470 nm) of RuBi-Glutamate (30 μM) before and during application of the glutamate transport blocker, TBOA (30 μM), in the (i) prelimbic or (ii) primary motor cortex of chow and extended access rats. Scale bar: 25 pA, 20 ms.

*(**F**)* Bar graph quantifying the amplitude of flash-evoked GLT-1 currents in astrocytes. GLT-1 current amplitude is not significantly different in (i) prelimbic cortex astrocytes from chow (n = 8/4) or extended (n = 7/4) access rats (t_(13)_ = 0.091, P = 0.929) or (ii) motor cortex astrocytes from chow (n = 7/4) or extended (n = 7/4) access rats. Bars represent mean ± s.e.m with individual values overlaid.

**Figure S2. Glutamatergic synapses are sensitive to cannabinoid modulation, but an endocannabinoid tone is not present in Extended access rats.**

(**A**) Example traces of evoked EPSCs before (black) and during (red) application of the CB1 receptor agonist, WIN55,212-2 (1 µM) in lOFC neurons from chow access rats.

(**B**) Time plot of evoked EPSC amplitude before and during application of WIN55,212-2 (1 µM) onto lOFC pyramidal neurons in Chow rats, n = 13/5.

(**C**) Bar graph quantifying the percentage inhibition of evoked AMPA EPSCs by WIN55,212-2 (1 µM) in chow access rats. WIN55,212-2 inhibits evoked EPSCs to a maximum of 77 ± 4%. This effect is significantly different from the baseline of 100% (t_(12)_ = 5.653, P = 0.0001, n = 13/5).

(**D**) Example traces of evoked EPSCs before (black) and during (red) application of the neutral CB1 receptor antagonist, NESS-0327 (0.5 µM) in lOFC pyramidal neurons from extended access rats.

(**E**) Time plot of evoked EPSC amplitude before and during application of NESS-0327 (0.5 µM) in lOFC neurons from extended access rats n = 7/4.

(**F**) Bar graph quantifying the percentage change in evoked AMPA EPSCs by NESS-0327 (0.5 µM) in extended access rats. NESS-0327 does not significantly alter evoked EPSC amplitude from the baseline of 100% (t_(6)_ = 0.404, P = 0.7001, n = 7/4). Bars represent mean ± s.e.m with individual values overlaid.

**Figure S3. GABAergic synapses onto lOFC pyramidal neurons exhibit endocannabinoid-induced long-term depression.**

(**A**) Experimental schematic illustrating theta-burst stimulation (TBS) of GABA inputs onto lOFC pyramidal neurons in the presence or absence of the CB1 receptor antagonist, AM251 or the mGluR5 antagonist, MTEP.

(**B**) Example traces of evoked IPSCs in lOFC neurons of naïve rats before (black) and after (blue) TBS from slices that were untreated (control) or treated with AM251 (3 µM) or MTEP (10 µM) onto lOFC neurons of naïve rats. Scale bars: 250 pA, 25 ms.

(**C**) Time plot of evoked IPSC amplitude before and after TBS in lOFC neurons of naïve rats (n = 8/4). TBS was blocked by AM251 (3 µM, n = 17/5) or MTEP (10 µM, n = 8/4). Dotted vertical lines indicate the 25-30 min time period following TBS analyzed in (D). There was a significant interaction of drug group x timecourse (F(96,1174) = 4.752, P <0.0001) and a main effect of drug (F(2,28) = 35.48, P<0.0001**** (Mixed effects model 2-way RM ANOVA).

(**D**) Bar graph quantifying the percentage inhibition of evoked IPSC amplitude 25-30 min following TBS in lOFC neurons from naive rats. TBS-LTD is present in lOFC neurons (n = 8/4), but not AM251- (n = 11/5) or MTEP-treated lOFC neurons (n = 7/4), One-way ANOVA: (F(2,23)=31.92, P <0.0001), Tukey’s multiple comparison test control vs AM251, P < 0.0001^####^, control vs MTEP, P < 0.0001^####^, AM251 vs MTEP, P > 0.05. Evoked IPSC amplitude after TBS is significantly different from baseline in lOFC neurons from control (t_(7)_ = 7.186, P=0.0002***), AM251-treated (t_(10)_ = 3.139, P=0.0105**), but not MTEP-treated neurons (t_(6)_ = 0.7110, P=0.504).

(**E**) Bar graph of the paired-pulse ratio of evoked IPSC amplitude before (Baseline) and after TBS (Post-TBS) in control, AM251-treated, or MTEP-treated lOFC neurons. A RM 2-way ANOVA gives a significant drug x LTD interaction (F(2, 20) = 5.804, P = 0.0103). A Sidak’s multiple comparison test reveals a significant effect of TBS on paired-pulse facilitation in control lOFC neurons (n = 8/4, P < 0.05^#^), but not AM251 (n = 8/5, P > 0.05) or MTEP-treated neurons (n = 6/4, P > 0.05).

Bars represent mean ± s.e.m with individual values overlaid.

**FIGURE S4. Electrophysiological properties of astrocytes are not different between chow and extended access rats.**

(**A**) Example traces from SR-101-labelled cells recorded in the (i) lOFC, (ii) prelimbic or (iii) primary motor cortex of chow and extended access rats.

(**B**) Current-voltage relationships from SR-101-labelled cells recorded in the (i) lOFC (n/N = 28/9 and 19/5), (ii) prelimbic (n/N = 8/5 and 8/4) and (iii) motor cortex (n/N = 8/5 and 7/4) of chow and extended access rats, respectively. There was a passive membrane conductance and lack of action potential firing indicative of astrocytes.

(**C**) Bar graph quantifying the resting membrane potential in SR-101-labelled cells from the (i) lOFC (n/N = 28/9 and 19/5), (ii) prelimbic (n/N = 8/5 and 8/4) and (iii) motor cortex (n/N = 8/5 and 7/4) of chow and extended access rats, respectively. There are no significant differences between groups in the lOFC (interaction: F(1,67) = 0.6011, P = 0.4409, diet effect: F(1,67) = 0.7696, P = 0.3835, drug effect: F(1,67) = 0.1839, P = 0.6694), mPFC (t(14) = 0.053, P = 0.95) or motor cortex (t(12) = 0.226, P = 0.8248).

(**D**) Bar graph quantifying the input resistance in SR-101-labelled cells from the (i) lOFC (n/N = 28/9 and 19/5), (ii) prelimbic (n/N = 8/5 and 8/4), and (iii) motor cortex (n/N = 8/5 and 7/4) of chow and extended access rats, respectively. There are no significant differences between groups in the lOFC (interaction: F(1,63) = 3.009, P = 0.0877, diet effect: F(1,63) = 0.7843, P = 0.3792, drug effect: F(1,63) = 0.5200, P = 0.4735), mPFC (t(14) = 1.143, P = 0.27) or motor cortex (t(12) = 0.698, P = 0.4984).

## Star Methods

### Subjects

All experiments on rats were approved by the Animal Care Committee of the University of Calgary, under guidelines set by the Canadian Council on Animal Care. Male Long-Evans rats (P50-55) were obtained from Charles River Laboratories and were individually housed in cages with paper bedding (Alphapads, Lawrenceville, GA) on a 12:12 reverse light dark cycle (lights on at 8:00 am).

### Diets

In addition to ad-libitum chow and water, rats were given access to a cafeteria diet (Kraft^TM^ smooth peanut butter, chocolate Timbit^TM^ donut holes, Kirkland^TM^ beef hotdogs, Froot Loops^TM^, Doritos^TM^) for either 24 h per day (Extended access), 1 h per day (Restricted access) or 0 h per day (Chow only) for 40 consecutive days. Energy density and macronutrient composition is listed in supplemental Table 1. Rats with restricted access to the cafeteria diet were given their 1 h access to cafeteria diet approximately 2 h into their dark cycle in addition to 24h access to chow. This restricted access group served as a pair fed control, in which animals had access to the cafeteria diet, but their ability to consume the diet was limited by a short duration of exposure (Ellacott et al., 2010). Rats’ body weights were measured immediately before and after 40 days of exposure to the cafeteria diet.

### Slice preparation

Rats were deeply anaesthetized with isoflurane and intracardially perfused with N-methyl D-glucamine (NMDG) solution of composition (in mM): 93 NMDG, 2.5 KCl, 1.2 NaH_2_PO_4_.H_2_O, 30 NaHCO_3_, 20 HEPES, 25 D-glucose, 5 sodium ascorbate, 3 sodium pyruvate, 2 thiourea, 10 MgSO_4_.7H_2_O, 0.5 CaCl_2_.2H_2_O. Rats were then decapitated and coronal sections (300 µm) containing orbitofrontal cortex (OFC) were cut in NMDG solution using a vibratome (VT1200, Leica Microsystems, Nussloch, Germany). Slices were recovered in warm NMDG solution (32 °C) for 10-12 min before being transferred to a holding chamber containing artificial cerebrospinal fluid (ACSF) of composition (in mM): 126 NaCl, 1.6 KCl, 1.1 NaH_2_PO_4_, 1.4 MgCl_2_, 2.4 CaCl_2_, 26 NaHCO_3_, 11 glucose (32 °C); equilibrated with 95% O2 / 5% CO_2_.

### Neuronal Electrophysiology

Slices were transferred to a chamber on an upright microscope (Olympus BX51WI) and continuously superfused with ACSF (2 mL.min^-1^, 34 °C). OFC neurons were visualized with a 40X water immersion objective using Dodt gradient contrast optics. Whole-cell voltage-clamp recordings (holding potential = -70 mV) of synaptic currents were made using a MultiClamp 700B amplifier (Axon Instruments, Molecular Devices). Pyramidal neurons in layer II/III of the lateral OFC (∼ 100-300 µm above the inflection point of the rhinal sulcus) were identified by morphological and electrophysiological characteristics (triangular shape, capacitance > 100 pF, input resistance < 100 mOhm). For recording of inhibitory postsynaptic currents (IPSCs), recordings were obtained with an internal solution of the following composition (in mM): 80 CsCH_3_SO_3_, 60 CsCl, 10 HEPES, 0.2 EGTA, 1 MgCl_2_, 2 MgATP, 0.3 NaGTP, 5 QX-314-Cl (pH 7.2–7.3, 305 mOsm). Inhibitory postsynaptic currents (IPSCs) were pharmacologically isolated with the □-amino-3-hydroxy-5 methyl-4-isoxazolepropionic acid (AMPA)/kainate receptor antagonist, DNQX (10 μM) and the glycine receptor antagonist, strychnine (1 μM). For recording of excitatory postsynaptic currents (EPSCs), recordings were made using an internal solution containing (in mM): 139 CsCH_3_SO_3_, 8 CsCl, 10 HEPES, 0.25 EGTA, 4 MgATP, 0.3 NaGTP, 7 Na-phosphocreatine (pH 7.2–7.3, 305 mOsm). EPSCs were pharmacologically isolated with the GABA_A_ receptor antagonist, picrotoxin (100 μM) and strychnine (1 μM). Series resistance (5-20 MΩ) and input resistance were monitored online with a 10 mV voltage step given 400 ms every stimulus. IPSCs and EPSCs were filtered at 2 kHz, digitized at 10 kHz and collected using pCLAMP 10 software. In the majority of experiments, electrically-evoked currents were elicited using a bipolar tungsten stimulating electrode (FHC, Maine, USA) placed ∼ 50-300 µm laterally from the recorded neuron. Theta-burst stimulation (TBS) consisted of 8 theta-burst trains. Each train contained 10 bursts (200 ms interburst interval), with each burst consisting of 5 stimuli at 100 Hz. Example traces of evoked EPSCs or IPSCs were constructed from averages of 30 sweeps. In a subset of experiments, spontaneous miniature EPSCs or mIPSCs were recorded in the presence of the sodium channel blocker, tetrodotoxin (TTX, 0.5 μM) to inhibit action-potential dependent activity. Individual quantal events were selected based on amplitude (>10 pA), decay time (<10 ms) and rise time (< 4 ms) and analyzed using Minianalysis (Synaptosoft).

### Astrocyte electrophysiology

Following tissue slicing and recovery, slices were transferred to a temporary holding chamber containing ACSF at physiological temperature and treated with sulforhodamine-101 dye (SR101, 20 μM, 20 min) uptake by astrocytes. Slices were then transferred back to a regular ACSF holding solution for at least 30 min. For electrophysiology experiments, SR-101-labelled OFC astrocytes were patch-clamped with a glass electrode (∼ 4-5 mΩ) containing internal solution of the following composition (in mM): 130 K-Gluconate, 10 KCl, 10 HEPES, 0.5 EGTA, 10 Na-phosphocreatine, 4 MgATP, 0.3 NaGTP. In addition to SR-101 labelling, astrocytes were confirmed by their small soma size (∼ 10 μm), low resting membrane potential (< -80 mV), low input resistance (< 10 mΩ), passive membrane properties (linear current-voltage (IV) relationship) and a lack of action potential firing. These properties distinguish astrocytes from oligodendrocytes, which have a higher input resistance of ∼60 m∧ (Meyer et al., 2018). Astrocytes were recorded in voltage-clamp configuration (-80 mV) in the presence of the following synaptic blockers: DNQX (10 μM), picrotoxin (100 μM), strychnine (1 μM), the N-methyl-d-aspartate (NMDA) receptor antagonist, D-AP5 (50 μM) and the non-selective Group I/II metabotropic glutamate receptor antagonist, MCPG (250 μM). Glutamate was applied onto astrocytes by uncaging Rubi-glutamate via flash photolysis (473 nm, 2 ms, Thorlabs). At the conclusion of each recording, the glutamate transport blocker, TBOA (30 μM) was finally added to abolish the recorded current. Remaining residual currents were subtracted from the original response to specifically isolate a glutamate transporter current.

### Astrocyte Dye-filling

OFC astrocytes located at a depth of ∼ 20-50 μM from the slice surface were patch-clamped with an internal solution of the following composition (in mM): 108 K-gluconate, 8 Na-gluconate, 2 MgCl_2_, 10 HEPES, 1 K_2_-EGTA, 4 K_2_-ATP, and 0.3 Na_3_-GTP. Once a giga Ω seal was observed, whole-cell configuration was obtained to initiate filling of astrocytes with Dextran-FITC (3-5 kDa) dye. Access resistance was maintained below 20 M Ω throughout the experiment. Cells were allowed to fill for a minimum of 10 min before z-stack, allowing visualisation of the full morphology of individual astrocytes. Astrocytes were identified by their SR101 dye uptake, presence of vascular endfeet (observed following FITC-dextran loading) and low input resistance (10-20 MΩ). Fluorescence imaging was performed on a custom two-photon laser-scanning microscope optimized for acute brain slices and patch-clamp electrophysiology. The microscope was equipped with a Ti:Sapph laser (Ultra II, □4 W average power, 670–1080 nm, Coherent), objectives (Zeiss 40× NA 1.0, Nikon 16× NA 0.8), a green bandpass emission filter (525–40 nm), an orange/red bandpass emission filter (605–70 nm), and associated photomultiplier tubes (GaAsP Hamamatsu). *Z*-stacks were used to assess the extent of FITC-dextran (3-5kDa) diffusion in individual astrocytes. To image FITC-dextran during astrocyte filling, the Ti:Sapph laser was tuned to 800 nm.

### Western Blot Analysis

Western blot analysis for GLT-1 was performed on samples of OFC dissected from 100 µm thick frozen sections. Samples were homogenized and sonicated in ice□cold RIPA buffer (Thermo Scientific, 89900) with cOmplete Protease Inhibitor Cocktail (Roche. 11836170001) and PhosSTOP (Roche, 4906845001). The homogenate was centrifuged at 13,000 ***g*** for 20 min at 4°C. The supernatant was collected and protein content was measured using a Pierce BCA Protein Assay Kit (Thermo Scientific, 23225). The samples were denatured with a loading buffer containing 2-mercaptoethanol for 20 minutes at room temperate then the mixture (20 µg/lane) was loaded onto a 10% sodium dodecyl sulphate□polyacrylamide gel electrophoresis (PAGE) gel and run on a minigel system (Bio Rad., Mississauga, ON, Canada). After the protein was transferred to PVDF membrane following electrophoresis, the membrane was blocked in 5% milk followed by the incubation with rabbit anti GLT-1 (Cell Signaling, 3838) at 1:250 at 4°C for 24 hours. For loading control, mouse beta-actin (GenScript, A00702) at 1:2500 was applied at room temperature for 1 hour. The blots were further incubated at room temperature for 1 h with horseradish peroxidase conjugated goat anti□rabbit IgG 1:2500 for GLT-1 or goat anti-mouse IgG 1:2500 for beta-actin (Thermo Scientific, 31460 and 31430). Finally, antibody binding was visualized using the enhanced chemiluminescence (ECL) system (Thermo Scientific, 32109) and scanned with a Gel Doc system (Bio□Rad, Mississauga, ON, Canada). The band density of GLT-1 (MW 65 kDa) was analyzed with ImageJ and normalized against the loading control.

### Immunohistochemistry

Coronal sections containing the OFC were incubated with blocking solution (5% goat serum, 1% BSA, 0.5% Triton X-100 in 0.1 M PBS) to prevent nonspecific binding. Floating sections were incubated in primary antibody GFAP (1:1000; MAB360, Millipore) overnight at 4°C, washed, and then incubated with secondary antibody (goat anti-rabbit, 1: 500) labelled with Alexa Fluor 488 for 1 h at room temperature. Slices were then mounted on Superfrost Plus clear slides (VWR international), dried, coated with Fluromount medium (Diagnostic Biosystems, Pleasanton, CA, USA) and cover slipped for imaging. Confocal fluorescent images were collected using Nikon Eclipse C1si spectral confocal microscope with motorized stage (Nikon Canada Inc., Ontario, CANADA). The objectives used were 10x PLAN APO DIC (NA 0.45), 20X PLAN APO DIC (NA 0.75) and 60x PLAN APO water immersion DIC (NA 1.20). The laser used was centered at 488 nm (with 515/30 emission filter) wavelengths for detecting GFAP. Threshold levels were adjusted to 20 (min) to 225 (max) pixels and particle analysis was based on size restriction of 0 cm² to infinity. All the images were employed the same settings to be consistent for image analysis.

Four coronal slices from the OFC were taken from 5 rats per group. Within each slice, four non-overlapping images per brain region were imaged for a total of 16 images per brain region of a given animal. A total of 80 fields of view were captured per animal at 60x magnification for GFAP staining. All the images were taken using imaging software EZ C1 for Nikon Confocal, Silver Version 3.91 (Nikon, Canada Inc., Ontario, CANADA). Stacked images were acquired using two frames with a resolution of 512×512 pixels. Offline image processing included maximal intensity projections was conducted using NIS elements. Fluorescence intensity (intensity units, IU) and percentage area of fluorescence was quantified for each image using Image J (1.48v, Wayne Rasband, National Institute of Health, USA). All images were taken blinded to the treatment of the rats.

### Data analysis and Statistics

All statistical analyses were performed in GraphPad Prism 7.02 (GraphPad, US). A two-way repeated measures analysis of variance (RM-ANOVA) was used to examine effects of two factors between multiple groups. With timecourse data, some cells died before the end of recording and therefore matching on every point could not be achieved, therefore we used a mixed-effect model (REML) repeated measures 2-way ANOVA. A one-way ANOVA was used to compare the effect of a treatment between multiple groups. Unless otherwise indicated, a Tukey’s multiple comparison test was used to assess within group differences. A two-sample t-test was used to compare between two groups when normality distribution was assumed. A one sampled t-test was used to assess difference from 100%. Significance was set at P<0.05. A Shapiro-Wilk test was used to assess for normal distribution of data. All data are expressed as mean ± standard error of the mean (SEM). Individual responses are plotted over averaged responses. Experimental designs and samples sizes aimed at minimizing usage and distress of rats, and were sufficient for detecting robust effect sizes. N refers to number of rats whereas n refers to number of cells/number of rats.

---

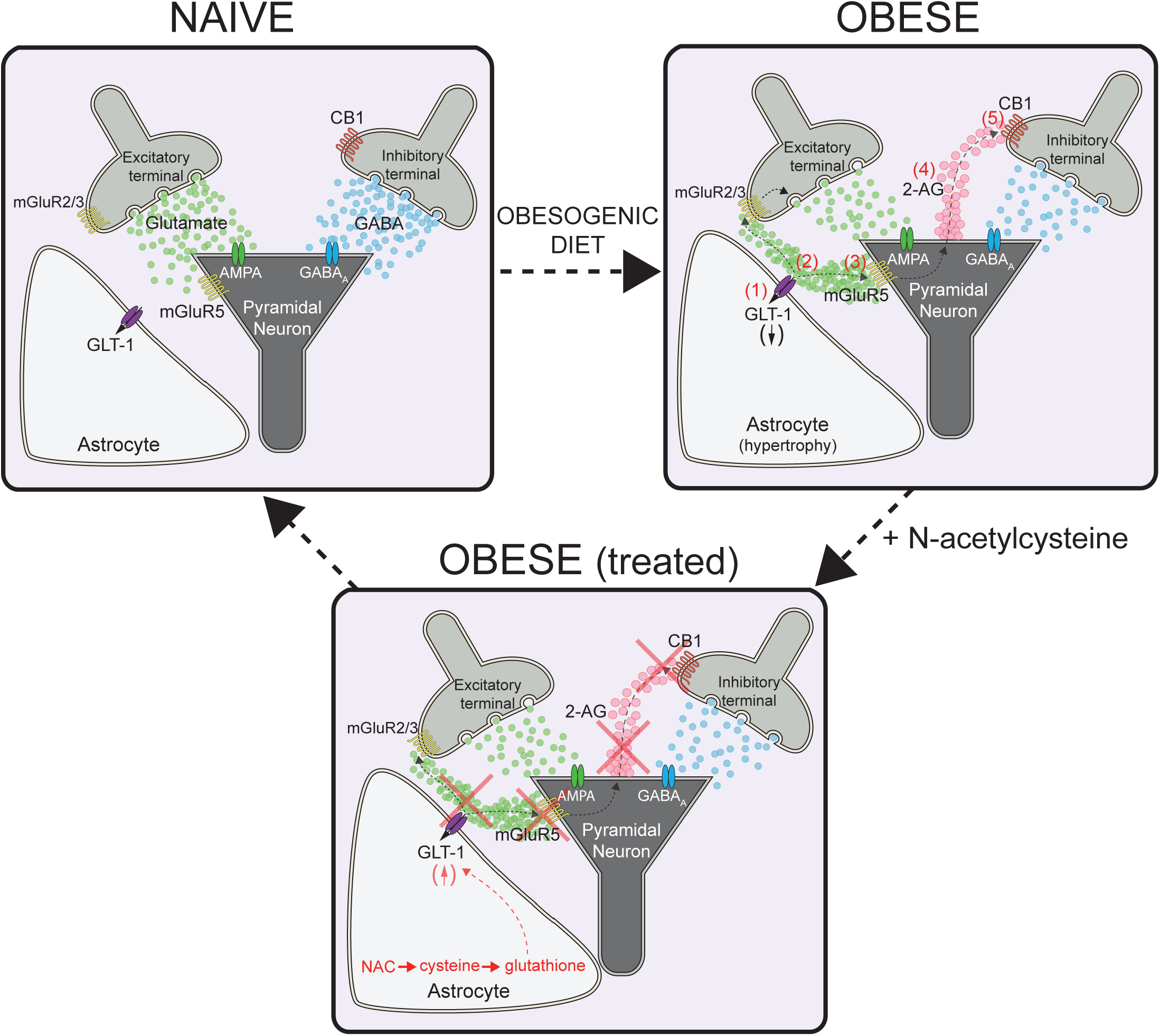

## Notes

### Competing Interest Statement

The authors have declared no competing interest.

### Summary of Updates

We have added new experiments and rewritten sections of the manuscript to improve testing of the hypothesis. Substantial changes are to figure 7 and supplemental figures 1-4.

