## Supplementary figures and images for "Obesity-induced astrocyte dysfunction impairs heterosynaptic plasticity in the orbitofrontal cortex"

### Supplemental Figures

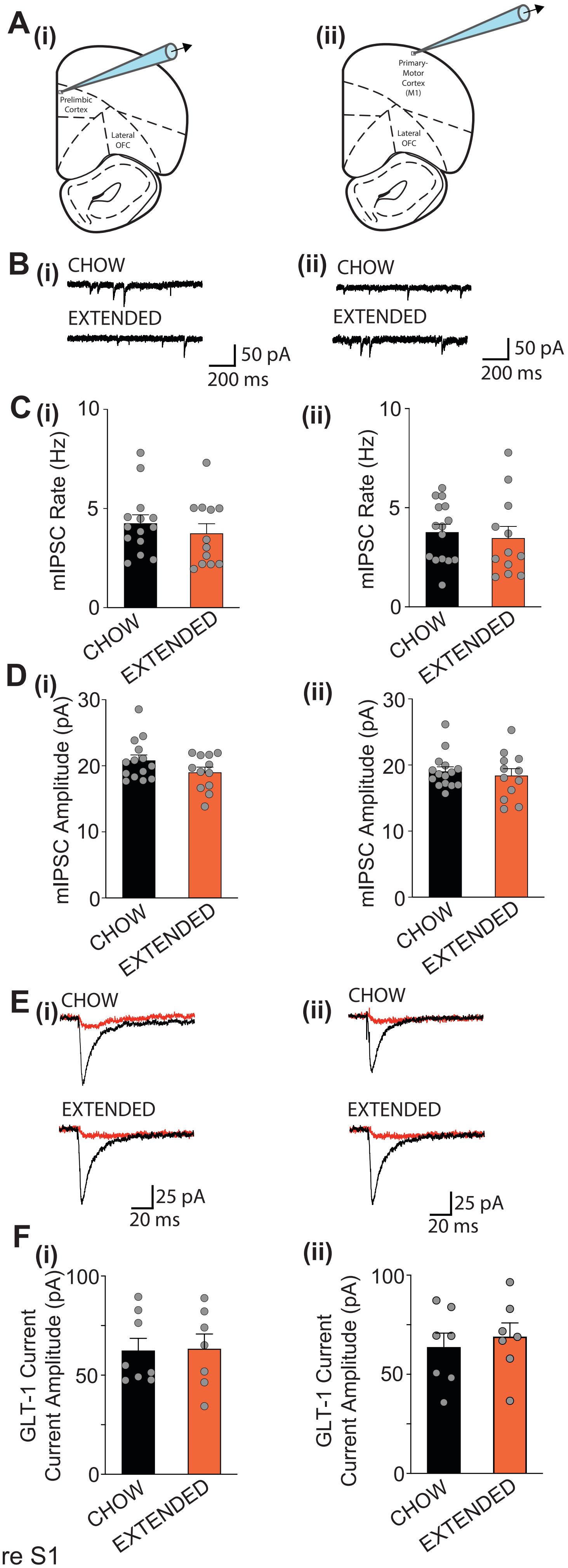

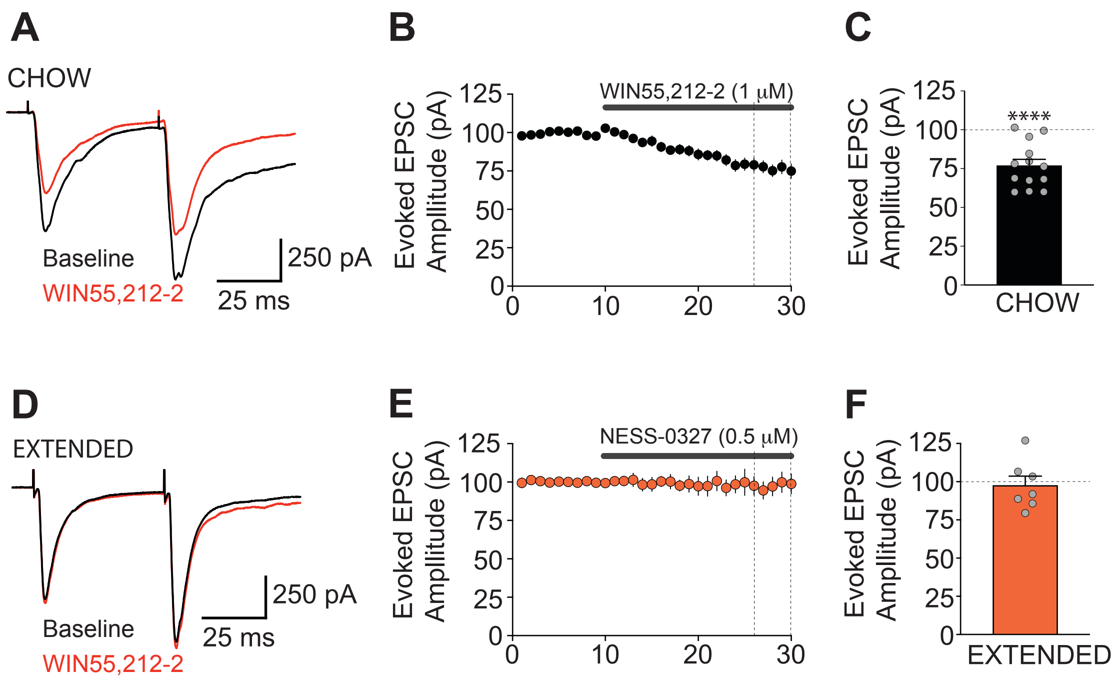

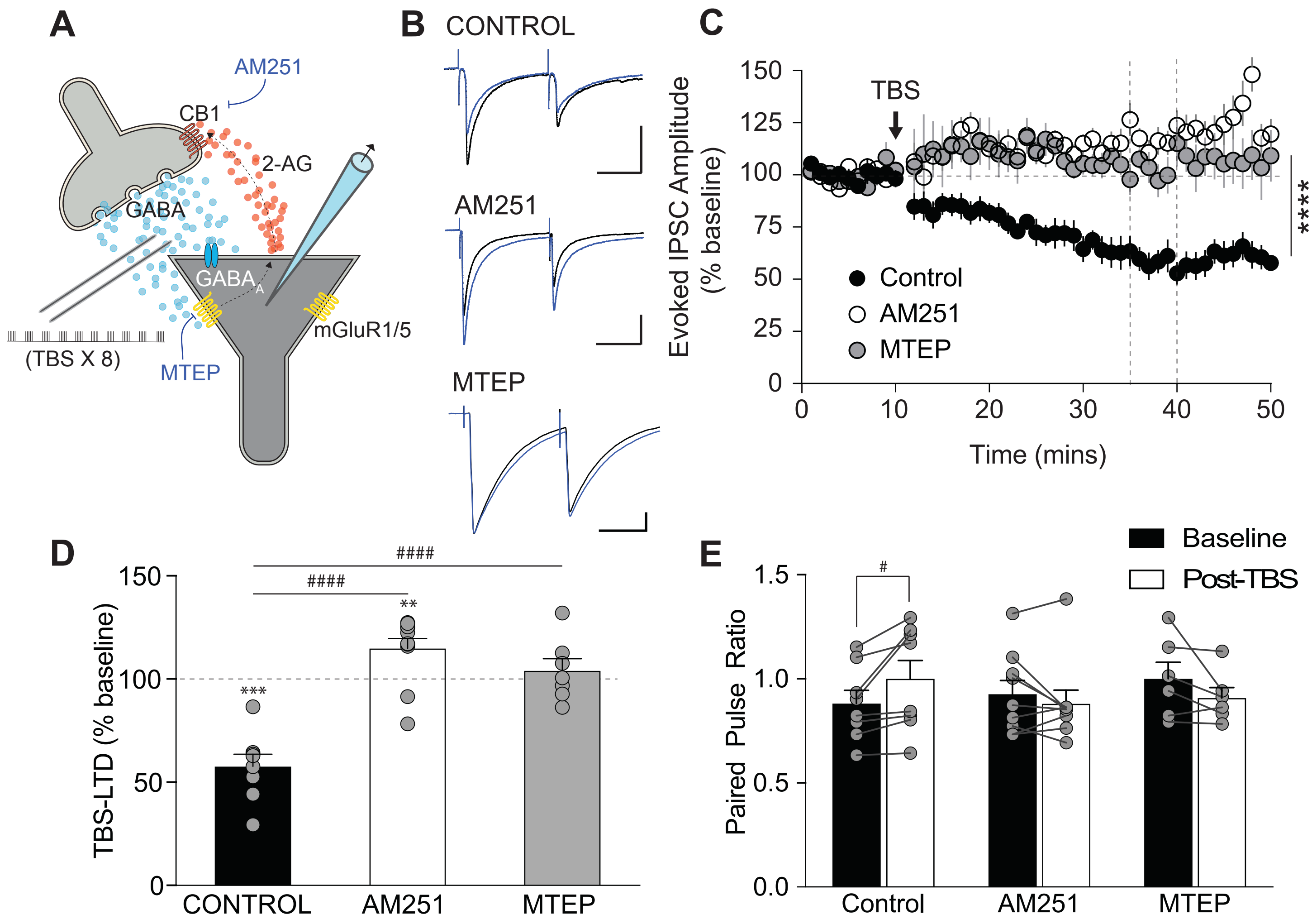

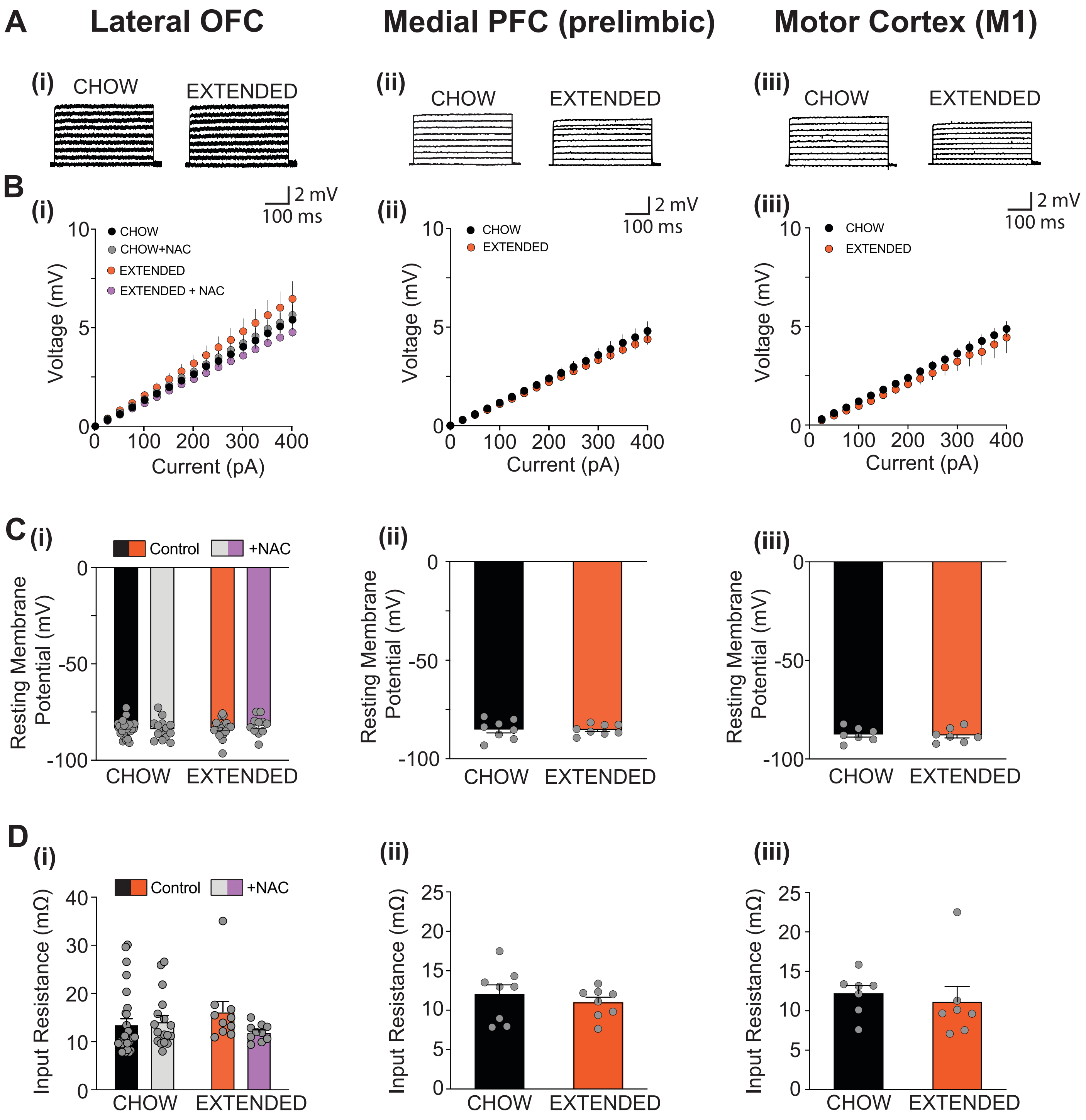
